# Phylogenetic coherence in microbiome composition across environmental gradients

**DOI:** 10.64898/2026.06.07.730742

**Authors:** Milena S. Chakraverti-Wuerthwein, Alissa Domenig, Seppe Kuehn

**Author notes:** Corresponding author: Seppe Kuehn, Department of Ecology and Evolution, University of Chicago.

## Abstract

Global surveys of microbial communities across biomes have shown that environmental variables such as depth and pH are strong determinants of community composition. However, we do not understand how the traits of individual taxa, and their evolutionary conservation, conspire to give rise to these patterns. Exploiting large-scale surveys of top soil and marine microbiomes, we use canonical correlation analysis (CCA) to concurrently infer directions of environmental variation and the associated compositional changes. We find that the primary canonical direction, capturing the dominant environmental gradient, exhibits a strong phylogenetic signal: individual species’ responses to environmental shifts along this direction are similar among taxa with shared evolutionary history. In contrast, secondary canonical directions show weak or no phylogenetic structure. Together, these results suggest a two-scale view of microbial community assembly. Deeply evolutionarily conserved traits govern community reorganization along the main environmental driver of community composition. Additional environmentally driven changes in community composition then reflect traits that are more evolutionarily labile.

## INTRODUCTION

A longstanding view in microbial ecology is that sufficient dispersal gives rise to community composition primarily determined by local environmental conditions. Global surveys across a variety of biomes have repeatedly demonstrated consistent associations between community composition and environmental variables such as pH, temperature, salinity, and depth [1–9]. This effect, also known as environmental filtering, is captured by the hypothesis of Baas-Becking [10], commonly paraphrased as “everything is everywhere; the environment selects.” Given environmental filtering is likely acting through traits impacting the capacity of a given taxon to grow, persist, or compete under particular conditions, these patterns raise a deeper question about which traits are being selected and how broadly they are shared across the tree of life.

Traits have been hypothesized to vary in the degree to which they are evolutionarily conserved [11, 12]. Whether an environmental gradient selects on traits that are conserved versus more randomly dispersed on the tree could determine whether or not communities are phylogenetically clustered. Selection on conserved traits would cause closely-related taxa to respond in similar ways to an environmental gradient (clustering), while selection on more randomly dispersed traits would show no phylogenetic signal in the community response (Fig. 1). The existing literature has examined this question through the language of over- and underdispersion, asking whether evolutionarily similar lineages occupy similar environments more or less than expected and how that reflects a balance between environmental filtering and competitive exclusion [13–16]. This approach considers co-occurence of related species across samples without explicit consideration for how the environment changes between those samples. However, specific traits likely depend on particular environmental gradients, such as photosynthesis and the availability of light. Studying phylogenetic dispersion without considering these gradients risks confounding the phylogenetic signal from multiple traits.

**FIG. 1.**
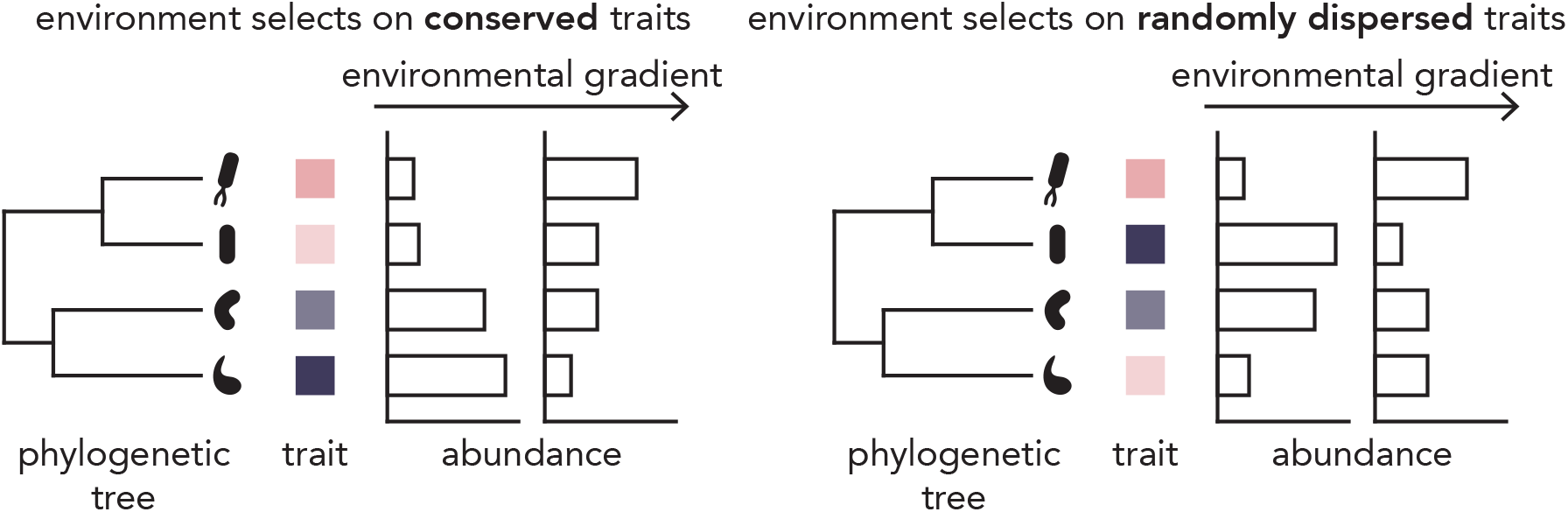
Phylogenetically coherent responses emerge for the dominant environment-composition association in global microbiomes. The primary environmental axis (left) selects on evolutionarily conserved traits, where closely related species (shown in a four-species, two-clade phylogeny) share similar trait values (pink to purple gradient) and exhibit similar abundance responses to environmental change. Secondary environmental axes (right) select on randomly dispersed traits, where trait values are phylogenetically independent, causing random abundance responses among closely related species.

One approach for explicitly considering environmental gradients is to correlate environmental variables such as pH or light with dominant axes of variation in community composition. In this case, the dominant axes are first identified using dimension reduction methods such as PCA, before being correlated with individual environmental variables [2, 5, 17]. This approach does not explicitly consider phylogeny and considers only a single environmental variable at a time. In reality, community composition does not vary independently with each variable we measure (e.g. pH), but instead changes jointly with multiple, sometimes correlated, environmental variables (pH, carbon, nitrogen etc) [18–20]. To understand how environment shapes community composition, and how this depends on evolutionary history we would like to learn axes of compositional and environmental change concurrently, across phylogenetic levels.

Here we use canonical correlation analysis to learn coupled axes of environmental and compositional variation. Each axis, or canonical direction, defines a gradient along which movement in environmental space corresponds to a correlated shift in community composition. By examining the phylogenetic structure of community response along each axis, we ask whether organisms with shared evolutionary lineages respond coherently to a given environmental gradient and whether this coherence differs between dominant and secondary axes of environmental variation.

We find that the primary axis, capturing the dominant mode of environment-composition covariation, exhibits strong phylogenetic structure: taxa within the same evolutionary lineages shift together in similar ways along this gradient. In contrast, secondary axes show less phylogenetic coherence. These results support a refined view of Baas-Becking’s hypothesis: the dominant environmental driver of community composition acts through deeply conserved traits, producing predictable, phylogenetically structured responses, while additional environment-driven variation reflects traits that are more evolutionarily labile and therefore idiosyncratic across lineages (Fig. 1).

## RESULTS

### A. Identifying axes of environment-composition covariation

Using published global survey datasets [2, 4], we asked how taxonomic composition covaries with measured environmental conditions on a global scale. We focused on datasets with substantial sample sizes (soil: *N* = 217; ocean: *N* = 125) and paired measurements of bacterial relative abundances via 16S amplicon sequencing and environmental variables (Fig. 2a). For the soil dataset, the environmental variables were: pH, total organic carbon (C), total nitrogen (N), total phosphorus (P), and soil texture (clay-silt, percent of fine fractions), for the ocean dataset, the environmental variables were: depth, temperature, phosphate concentration (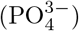), nitrite concentration (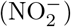), salinity, and dissolved oxygen (O_2_). These variables were selected from a broader set to reduce highly collinear or sparsely measured variables (see SI Figs. S1 S2 S3 and Sec. S2 for more details on how the datasets were curated). This pairing allows us to relate changes in community composition to the environmental context in which communities are observed.

**FIG. 2.**
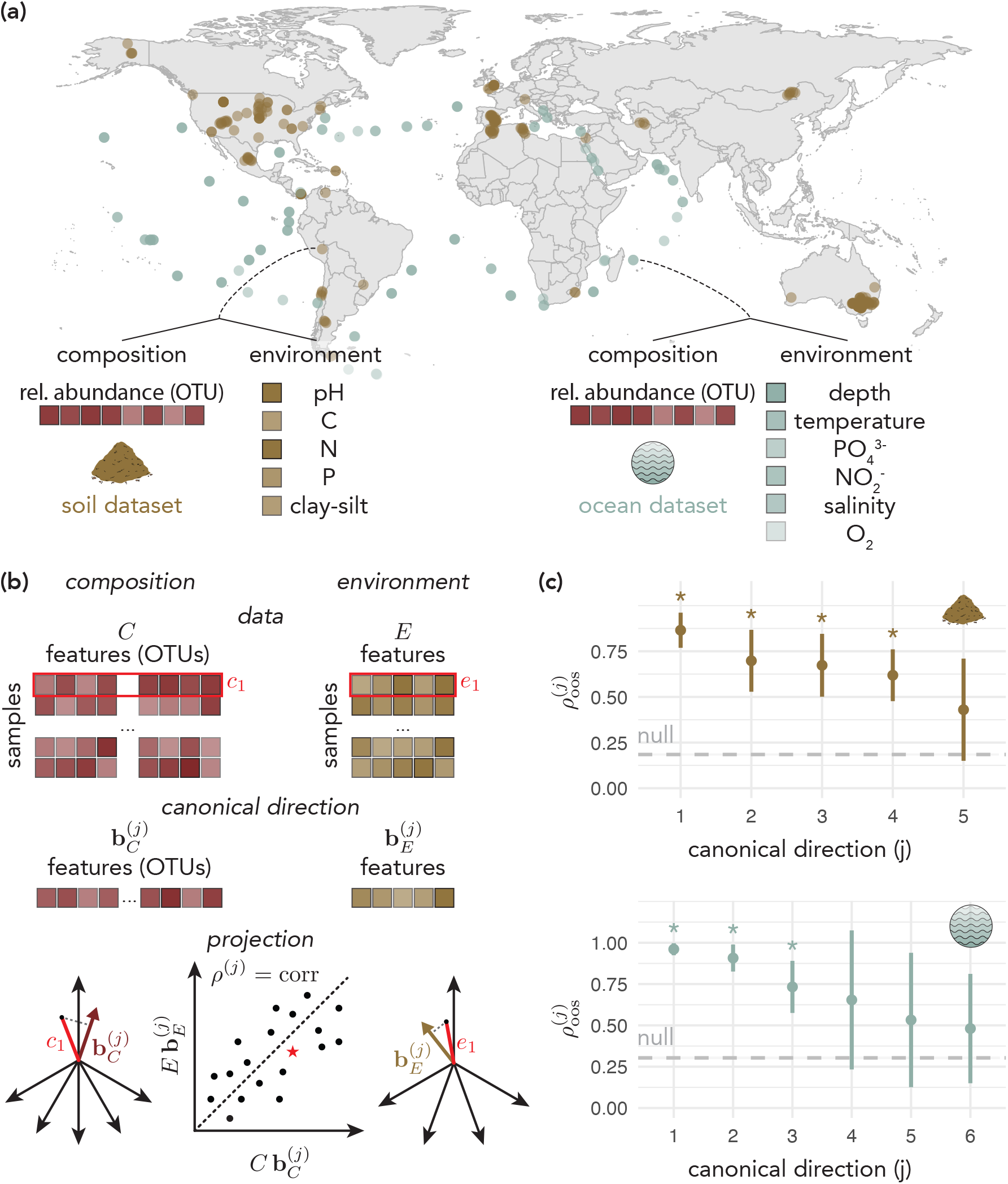
Canonical correlation analysis (CCA) identifies coupled gradients linking environment and community composition. **(a)** Global surveys pair microbial community profiles with environmental measurements across samples. The map shows sampling locations for soil (brown) [4] and ocean (blue) [2] datasets (points have fixed opacity, darker regions reflect overlapping locations of samples). Schematics indicate that each sample provides multivariate environmental context and a community profile that can be summarized by the relative abundances of different OTUs (opacity of the small squares is a schematic sketch to indicate different values for each measurement). **(b)** CCA finds paired canonical directions, a compositional loading vector, 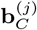, and an environmental loading vector, 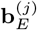, that maximize the correlation between the projected samples. **(c)** Canonical directions are evaluated against a shuffle-based null in which sample labels are permuted to destroy any environment-composition coupling; the null out-of-sample correlation shown in grey is the strongest (first) canonical direction attainable under permutation (see SI Sec. S3.B.4 for more details). 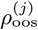 denotes the average out-of-sample correlation across *k* = 15 cross-validation folds (points show the mean *±* SD). For each canonical direction, we calculate *p* as the fraction of folds whose out-of-sample correlation falls at or below the null; asterisks mark directions for which every fold exceeds the null (*p <* 1*/k ≈* 0.067), namely CD1-4 for soil and CD1-3 for ocean (full values can be found in SI Table S1).

To quantify environment-composition coupling, we applied canonical correlation analysis (CCA), a linear dimensionality reduction technique that identifies paired gradients across environmental and compositional space. Unlike PCA, which finds directions of maximum variance within a single dataset, CCA learns directions that maximize covariation between two datasets. Specifically, CCA identifies pairs of basis vectors 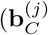 in compositional space and 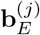 in environmental space) such that projections of samples onto these vectors yield maximally correlated coordinates (Fig. 2b). The loadings on these basis vectors reveal which environmental conditions are varying and which taxa are responding. We performed CCA on both soil and marine datasets.

In the global soil microbiome we identify 4 significant canonical directions (out of 5), whereas in the global ocean microbiome we identify 3 significant canonical directions (out of 6) (Fig. 2c). Significance was assessed by comparing out-of-sample correlations, 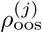, against a shuffle-based null that destroys environment-composition coupling, using the strongest (first) canonical direction obtained under permutation as a common reference for all true canonical directions (see SI Sec. S3.B.4, SI Fig. S9 for more details). We implemented regularized CCA [21] with *k*-fold (*k* = 15) cross-validation to learn the hyperparameter value and ensure robustness of the identified directions (for more details, see SI Sec. S3.B, SI Fig. S7).

### B. Relation to other methods

Several approaches have been used to link microbial community composition to environmental gradients and phylogeny. One common strategy first decomposes composition via PCA or PCoA and then correlates the leading components with individual environmental variables one at a time [2, 5, 17]; a second asks whether community composition is phylogenetically over- or under-dispersed across samples, without reference to specific measured environmental variables [13–16]. Both consider environmental drivers of composition one variable, or one undifferentiated axis, at a time, whereas community composition typically responds jointly to several correlated environmental factors (SI Fig. S1, S3). In practice, CCA and PCA recover the same dominant axis of compositional variation; however, while PCA’s leading components explain more variance in-sample, CCA’s canonical directions are more stable between training and held-out folds (SI Figs. S11, S12). See SI Sec. S3.C for a more in-depth review of these related approaches.

The approach most closely related to our work correlates each taxon’s abundance with an environmental axis and tests the phylogenetic conservation of the resulting “trait” (see, e.g., [22], where single environmental variables are used as the axes and consenTRAIT as the measure of phylogenetic conservation). When the axis is instead taken to be one of the environmental directions identified by CCA, this correlation-based trait is linearly related to our canonical weight trait, as shown in SI Sec. S3.C. The correlation captures each taxon’s total association with the environmental axis, including any it inherits through abundance covariance with other taxa, whereas the canonical weight reflects the taxon’s contribution after accounting for covariance, to a degree set by the regularization strength (with strong regularization, the two coincide up to a positive rescaling).

Thus, CCA provides a more holistic picture of environmental axes composed of multiple factors and their mapping to correlated changes in community composition.

### C. Dominant environmental gradients elicit phylogenetically coherent community responses

Having established significant environment-composition coupling, and the relevance of CCA to identifying this coupling, we next asked whether community responses are phylogenetically structured: do closely related taxa respond similarly across environmental gradients? Since taxonomic ranks correspond to nested clades of increasing phylogenetic depth, if responses are phylogenetically coherent, then the coupling should remain stable when composition is coarse-grained to higher taxonomic levels. To test this, we first performed closed reference mapping of ASVs (soils) or OTUs (ocean) to the GreenGenes2 phylogenetic tree (SI Figs. S4, S5, S6; see SI Sec. S2.C.2 for more details). For both datasets, we summed OTU relative abundances within each taxonomic level (genus, family, etc.) and repeated CCA using these coarse-grained abundances while holding environmental variables fixed (Fig. 3a). This aggregation approach offers a coarse, rank-based complement to the more classical, continuous coherence metrics described below (e.g. consenTRAIT [11] and Pagel’s *λ* [23]).

**FIG. 3.**
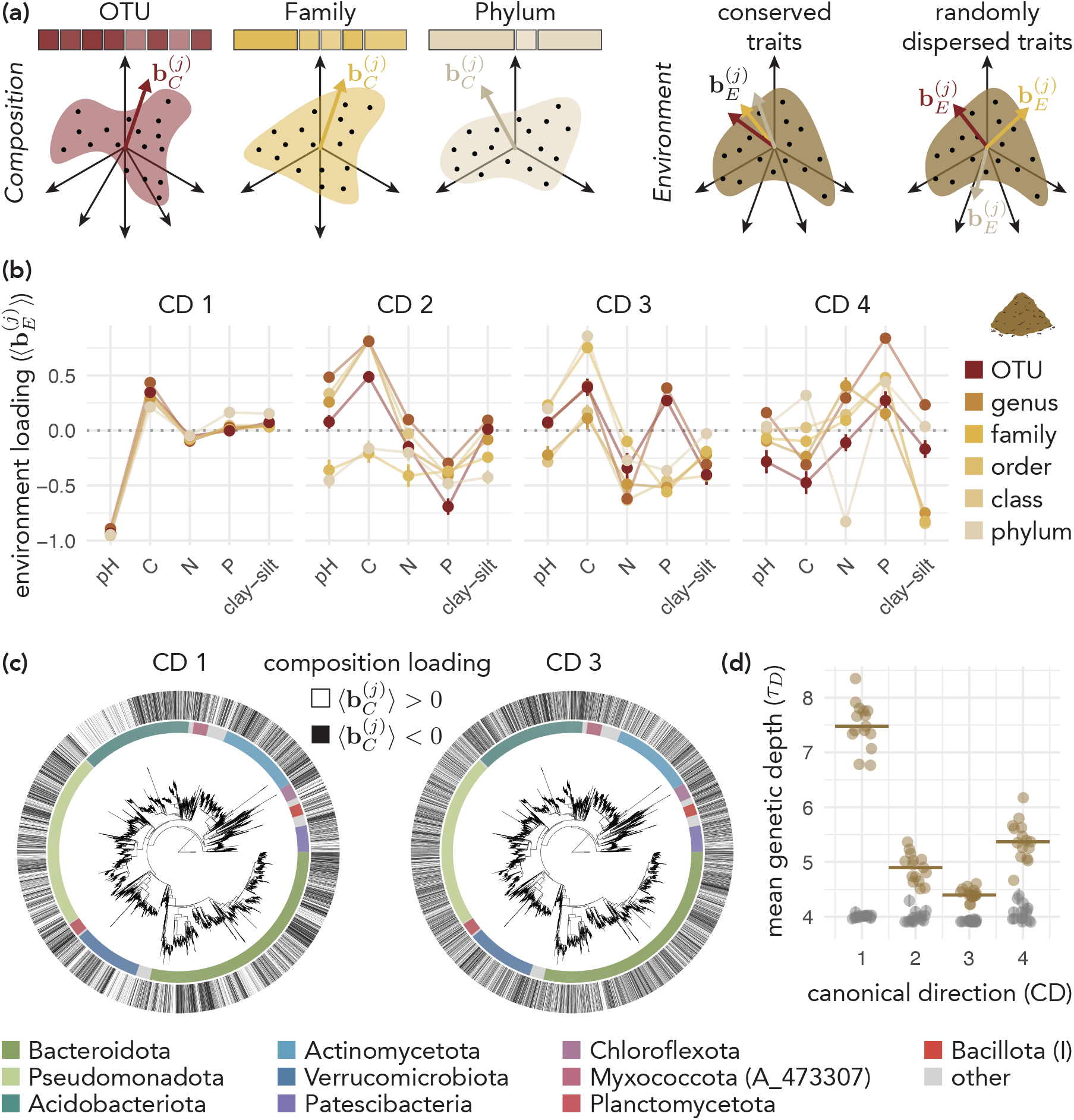
The dominant soil environment-composition axis (pH) is conserved across taxonomic levels and is phylogenetically coherent. **(a)** Schematic showing that phylogenetically coherent responses yield consistent environmental loading vectors across taxonomic resolutions, as closely related taxa with similar responses are aggregated together during coarse-graining. **(b)** The first canonical direction (CD1) in environmental space is consistently oriented across taxonomic resolutions, whereas higher-order directions (CD2–4) are less stable. Within each fold, canonical directions are sign-aligned to a common orientation before computing the mean *±* SD across CV folds (SI Fig. S8). **(c)** Community responses along CD1 exhibit clade-level structure. In other words, OTUs with positive/negative CD1 loadings cluster within phylogenetic lineages as qualitatively shown, while this pattern is weaker for CD2. OTU loadings are averaged across CV folds and then binarized by sign for display. Phyla are ordered by mean relative abundance across samples; the 10 most abundant are colored and the remainder are shown in grey. **(d)** Signed, size-weighted consenTRAIT *τ*_*D*_ for each canonical direction (for details, see SI Sec. S4.A). Here larger *τ*_*D*_ indicates that OTUs sharing the same loading sign cluster within deeper phylogenetic lineages than expected from a random permutation of loading signs. Each point gives *τ*_*D*_ for one canonical direction *j* and cross-validation fold index *f* ; horizontal bars show the median across folds. The primary canonical direction (CD1) shows deeper same-sign clades than subsequent directions.

In the soil dataset, the first canonical direction (CD1) in environmental space is strongly conserved across taxonomic levels and is aligned with pH, consistent with extensive work identifying pH as a major organizing axis of soil microbiome [17] (Fig. 3b, SI Figs. S13 and S14). In addition to the stability of the direction of CD1 in environmental space, the correlation strength is conserved across taxonomic levels. The out-of-sample canonical correlation for the first direction, 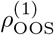, remains within 0.782*−*0.876 across all six taxonomic levels (SI Fig. S10). In contrast, the secondary canonical directions (CD2–4) are *incoherent* across taxonomic resolutions: the environmental gradients that CCA captures, and the specific environmental variables with highest loadings, shift depending on whether composition is represented at the OTU, genus, or phylum level. This instability indicates that these axes do not reflect phylogenetically coherent responses.

The same pattern is visible in the compositional loadings. Recall that CCA identifies basis vectors 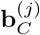 in compositional space (Fig. 2b); each OTU *α* has a loading 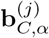 on canonical direction *j*. For the first canonical direction, mapping OTU loadings onto the phylogenetic tree reveals clades with the same sign, indicative of coherent responses to the corresponding environemntal direction. Whereas subsequent canonical directions show weaker phylogenetic clustering (Fig. 3c).

We quantified this observation using a modified version of consenTRAIT, a metric that measures phylogenetic conservation of microbial traits [11]. Here, we treat loadings of each OTU on each canonical direction as a “trait” and measure the phylogenetic conservation via the statistic *τ*_*D*_. The *τ*_*D*_ statistic measures the mean phylogenetic depth of clades in which at least 90% of OTUs share the same loading sign; larger *τ*_*D*_ indicates that same-sign OTUs cluster within deeper phylogenetic lineages (for more details see SI Sec. S4.A). To establish a null expectation in the absence of phylogenetic structure, we randomly permuted the loadings across the tips of the tree (*N* = 101 permutations) and recomputed *τ*_*D*_ for each permutation. The observed *τ*_*D*_ values (brown) exceed the null distribution (grey) for all canonical directions, confirming phylogenetic coherence, although coherence decays notably from the primary (CD1) to the secondary (CD2–4) directions (Fig. 3d). The depth of coherence in the primary direction corresponds to a phylogenetic depth of sub-genus-level classification, as benchmarked by calculating inter- and intra-group distances within the annotated reference tree (SI Fig. S15). We corroborated this finding with another standard metric of phylogenetic conservation, Pagel’s *λ* [23] (SI Sec. S4.C; SI Fig. S16).

Together, these results indicate that the dominant environment-composition axis is stable to taxonomic coarse-graining and corresponds to a phylogenetically conserved community response, whereas secondary axes are progressively less conserved.

### D. Ocean microbiomes exhibit the same phylogenetic structure despite different environmental drivers

This pattern of phylogenetically coherent primary axes and incoherent secondary axes is not unique to soils. In the ocean dataset, the primary canonical direction (CD1; Fig. 4a, SI Fig. S13, S14) and correlation strength (SI Fig. S10) is likewise conserved across taxonomic levels and is aligned with depth, consistent with prior work [2, 24]. Notably, while the dominant environmental gradient differs between ecosystems, pH in soils versus depth in the ocean, the phylogenetic structure of community responses is conserved. Secondary directions (CD2,3) again show incoherence across taxonomic levels, with environmental loadings shifting as composition is coarse-grained. Visual assessment of the OTU loadings on the tree (Fig. 4b), together with the adjusted consenTRAIT *τ*_*D*_ (Fig. 4c) and Pagel’s *λ* (SI Fig. S16), reveal a greater retention of phylogenetic signal in secondary directions for the ocean than in soils, but the same overall decay with canonical direction rank. Again, the degree of coherence in the primary direction corresponds to a phylogenetic depth of roughly genus-level classification (SI Fig. S15).

**FIG. 4.**
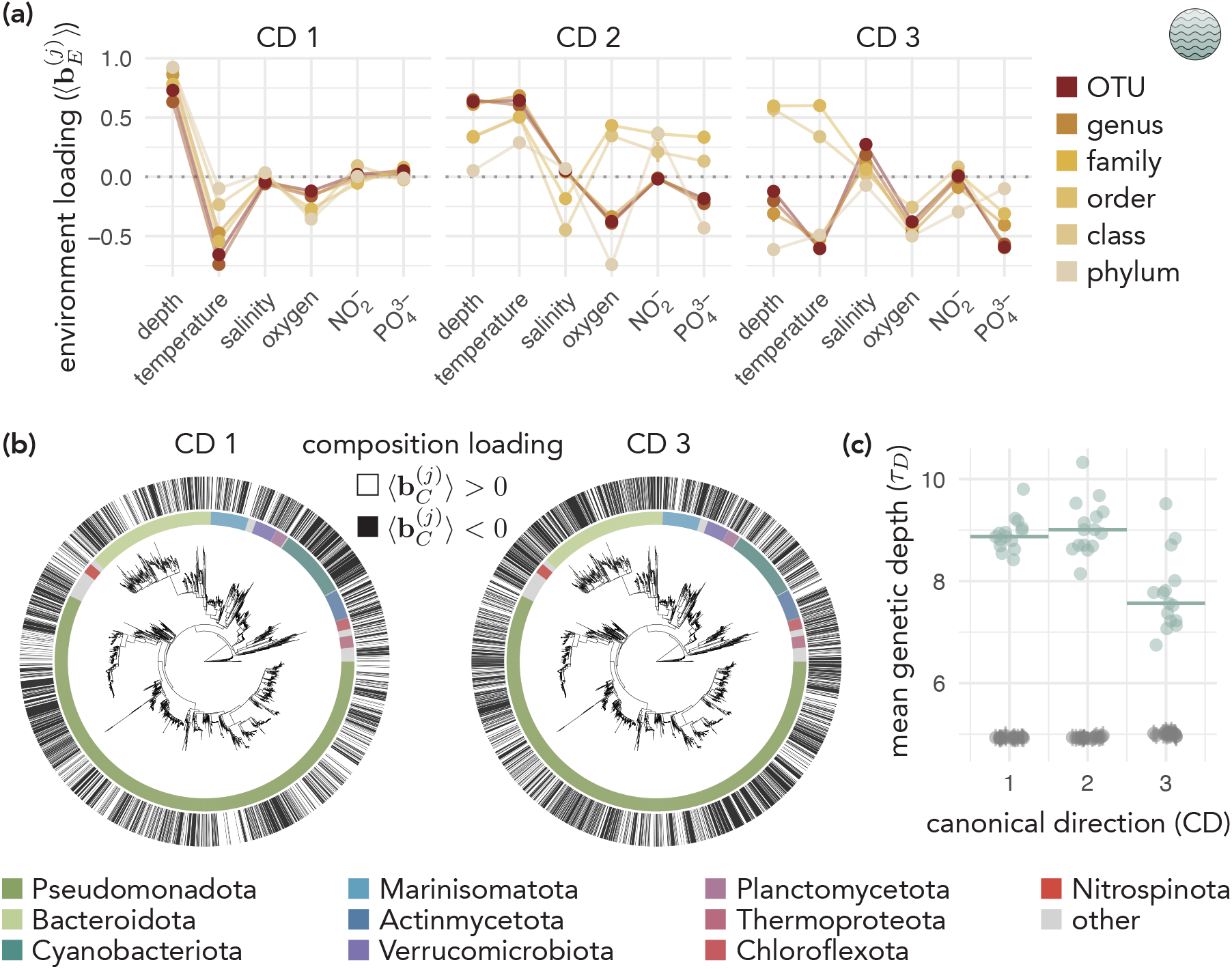
Phylogenetically coherent community responses to dominant environment-composition gradients extend beyond soils to the ocean microbiome. Format as in Fig. 3. **(a)** Reproducibility of environmental canonical directions across taxonomic levels (mean *±* SD across folds after sign alignment). **(b)** Phylogenetic distribution of OTU loading signs for CD1 and CD3, shown on the reference tree (coloring scheme as in Fig. 3). **(c)** Signed, size-weighted consenTRAIT *τ*_*D*_ for each canonical direction (see Fig. 3 for format and definitions); horizontal bars show the median across folds. The decay in phylogenetic coherence across canonical directions is shallower in oceans than in soils.

## DISCUSSION

Our results suggest a two-scale view of microbial community assembly: deeply evolutionarily conserved traits drive reorganization along the main environmental gradient, while more labile traits underlie additional environmentally driven variation. This pattern appears in both systems. In soils the primary axis is pH; in the ocean it is depth. Irrespective of the source of the gradient and the biome, the primary axis is stable when composition is aggregated to higher taxonomic ranks (Figs. 3b and 4a) and the OTUs that share a loading sign form clades that reach roughly sub-genus (soil) to genus (ocean) depth, which is deeper than expected under random permutation (Figs. 3d and 4c).

Our analysis of how environments structure communities rests on the premise that dispersal is sufficient for the pool of taxa available to selection to be similar across locations and environments, such that environmental factors are the main determinant of which taxa are abundant rather than distinct geographic effects. This assumption, often summarized as “everything is everywhere, but the environment selects” [10, 25], has been debated for a long time. Early arguments focused on protists and their large population sizes [26, 27], and large-scale surveys have since tried to separate the effects of environmental selection from those of dispersal [28, 29]. Many of these surveys support a primary role for environmental selection: bacterial composition in topsoil tracks environment, in particular pH, with little effect of distance [5, 17, 30]; environment explained more variation than distance in a synthesis of 54 studies across biomes [29]; and in the ocean, depth and temperature dominate while distance plays a subordinate role [2]. Dispersal limitation has nonetheless been documented, including in marine bacteria [31–33], and most studies detect some distance effect, more often at finer taxonomic resolution [28, 29]. This tension is hard to resolve with survey data, however, because dispersal has rarely been tested directly, and then only at small scales [34, 35]. Geographic distance is the usual proxy for limited dispersal, but it is entangled with unmeasured environmental variables that may covary with distance, so distance effects are potentially overestimated [29, 34].

Limited dispersal, however, need not undermine the argument that environment structures composition through traits conserved over long evolutionary history, provided the two processes operate on different timescales. In neutral simulations of ocean circulation, about 90% of the surface ocean is connected within a decade [36], and when neutral mutation is added, periodic takeovers between neighboring regions keep divergence between distant regions to about 0.5% of the genome, far below species-level differences [37]. Limited dispersal could therefore add regional structure among close relatives while maintaining a globally dispersed pool at deeper phylogenetic levels consistent with traits that have longer evolutionary history.

Regional variation at deeper clade levels, however, can still arise. For example, when thermal selection is added to the same ocean circulation model, populations do not cross intervening warmer or colder waters, and connectivity stays below 15% over a century [36]. In this case, dispersal limitation is itself a product of environmental selection, so the regional structure it creates would still reflect selection on environmentally relevant traits rather than confound it. There is also evidence that the geographic range sizes of taxa differ among bacterial phyla and families [38], indicating that dispersal ability may itself be a phylogenetically structured trait. Clades with limited dispersal would then be confined to particular regions. If the region a clade is confined to happens to lie at one end of the environmental gradient (for example, regions where soils tend to be alkaline), the clade could appear to respond coherently to that gradient even if the gradient does not select on its traits, inflating apparent coherence. If instead its confinement is unrelated to the gradient (for example, regions with mixed soil pH), the same environment could be occupied by different, possibly distantly related clades in different regions, adding variation unexplained by the measured environment and weakening apparent coherence. The fact that we observe coherence at sub-genus to genus depth indicates that dispersal limitation uncorrelated with the measured gradient does not erase the signal, but clade-level differences in dispersal that covary with the measured gradient remain a scenario our analysis cannot exclude.

Taking a step back, our results attempt to bridge a gap between how the environment structures communities via a selection on traits that evolution has constrained. We speculate that the depth of phylogenetic coherence along the primary environmental axis reflects selection on deep physiological commitments.

In soils, the primary environmental axis is pH, which constrains enzymatic function. Bacteria adapted to acidic or basic conditions seem to alter cytoplasmic pH tracking the external environment [39] in order to maintain a functional proton gradient [39, 40] (with a few known exceptions [41]). However, enzymes and proteins within the cell are generally pH sensitive due to local protonation states impacting function [42]. As a result, in addition to maintaining membrane mechanisms to cope with a particular external pH, the entirety of an organism’s internal enzyme repertoire must be tuned to the associated cytoplasmic pH. Adapting to substantial changes in pH would likely require the build up of many gradual evolutionary modifications and thus be deeply conserved, matching the coherent response of taxa in a community to a pH gradient.

In the ocean, the primary environmental axis is depth, which partitions fundamentally different energy metabolisms. Light is confined to the surface and the flux of organic carbon to depth declines as a power law [43], partitioning the water column into distinct energetic regimes: photoautotrophs in the euphotic zone producing the organic matter that sustains chemoheterotrophs below. Community composition is strongly stratified along this axis [2, 44]. The photoautotrophic lifestyle that underpins this structure is deeply conserved, with carbon fixation pathways and their associated machinery embedded in central metabolism [12, 45]. Unlike accessory traits exchanged readily via the mobilome, the transition between autotrophy and heterotrophy requires the coordinated reorganization of central carbon and energy metabolism. Just like in the case of pH adaptation, such deep metabolic commitments may be the underlying mechanism of the phylogenetic coherence observed across depth-stratified marine communities.

Connecting these physiological arguments back to the phylogenetic signal in environmental selection requires moving beyond inferring traits from taxonomy to measuring them directly. Systematic high-throughput phenotyping across phylogenetically diverse isolates could more directly test the framework developed here, linking measurable physiological traits to their co-varying environmental gradients and the phylogenetic depths at which they are conserved. Measurements of catabolic capabilities [46, 47] or pH optima [48] across a broad range of strains, would allow trait distributions to be mapped onto the phylogeny and compared against community-level patterns. More broadly, integrating high-throughput phenotyping with phylogenetically resolved community data offers a path toward mechanistically grounding the two-scale picture of environmental filtering proposed here.

Together, these results suggest a layered view of environmental filtering. Rather than treating the environment as a single selective force, we find that different environmental axes are associated with community responses at different depths of the phylogenetic tree. Thus, microbial community assembly is shaped not only by which environments taxa encounter, but by how the traits needed to persist in those environments are distributed across evolutionary history. This perspective reframes environmental filtering as an explicitly eco-evolutionary process: contemporary environments sort communities through traits whose accessibility is constrained by the evolutionary past.

## CODE AVAILABILITY

All code and pre-computed data used to generate the figures in this manuscript are deposited on GitHub (https://github.com/MilenaCW/microbiome-phylo-coherence) and archived on Zenodo (doi: 10.5281/zenodo.20586902).

## DATA AVAILABILITY

This study uses previously published, publicly available data. Soil 16S rRNA gene amplicon sequencing data and environmental metadata from Delgado-Baquerizo *et al*. [4] were obtained from Figshare (doi: 10.6084/m9.figshare.5611321). TARA Oceans miTAG OTU count tables and environmental metadata [2, 49] were obtained from https://ocean-microbiome.embl.de/companion.html, and carbonate chemistry metadata were obtained from PANGAEA [50]. The processed data used to generate all figures are available on Zenodo (doi: 10.5281/zenodo.20586902).

## AUTHOR CONTRIBUTIONS

M.S.C.-W. and S.K. designed research; M.S.C.-W. and A.D. performed research; M.S.C.-W. and A.D. analyzed data; and M.S.C.-W. and S.K. wrote the paper.

## DISCLOSURE OF DELEGATION TO GENERATIVE AI

The authors declare the use of generative AI in the research and writing process. According to the GAIDeT taxonomy (2025), the following tasks were delegated to GAI tools (Claude Sonnet, versions 4–5, and Claude Opus 5.5) under full human supervision: code generation; code optimization; text generation; proofreading and editing. Responsibility for the final manuscript lies entirely with the authors. GAI tools are not listed as authors and do not bear responsibility for the final outcomes.

## ACKNOWLEDGMENTS

We thank Arvind Murugan, Stefano Allesina, Akshit Goyal, Abby Skwara, as well as the Kuehn and Murugan groups at University of Chicago at large for helpful discussions. The authors acknowledge the University of Chicago’s Research Computing Center for computing resources, the Center for Living Systems (National Science Foundation award PHY-2317138), and the NSF-Simons National Institute for Mathematics and Theory in Biology (National Science Foundation award DMS-2235451 and Simons Foundation award MPS-NITMB-00005320). M.C.-W. acknowledges the Fannie and John Hertz Fellowship Award as well as the National Science Foundation Graduate Research Fellowship Program under grant number DGE 1746045. A.D. acknowledges the University of Chicago Quad Undergraduate Research Scholars Program. S.K. acknowledges the National Science Foundation CAREER Award under grant number BIO/MCB 2340416 and National Institute of General Medical Sciences under grant number R35GM164154. Any opinions, findings, conclusions or recommendations expressed in this material are those of the authors and do not necessarily reflect the views of the National Science Foundation.

