## Supplemental Information for "Phylogenetic coherence in microbiome composition across environmental gradients"

**CONTENTS**

S1. Code availability 4

S2. Data Curation 4

S2.A. Selection of Datasets 4

S2.A.1. Soil Dataset 4

S2.A.2. Ocean Dataset 4

S2.B. Preparing the Environmental Data 5

S2.B.1. Soil Dataset 5

S2.B.2. Ocean Dataset 5

S2.C. Preparing the Compositional Data 9

S2.C.1. DADA2 Processing (Soil Dataset) 9

S2.C.2. GreenGenes2 Assignment (Both Datasets) 9

S3. Canonical Correlation Analysis 12

S3.A. Technical Definitions 12

S3.A.1. Canonical Correlation Analysis 12

S3.A.2. Regularized CCA 14

S3.B. Computational Implementation 15

S3.B.1. Data Preprocessing 15

S3.B.2. $K$-fold Cross-Validation and Hyperparameter Search 15

S3.B.3. Loadings Extraction, Normalization, and Alignment 15

S3.B.4. Null Model 17

S3.B.5. CCA at Different Taxonomic Levels 19

S3.C. Relation to other methods 19

S3.C.1. PCA vs. CCA decomposition of community composition 19

S3.C.2. Correlation-based vs. canonical weight definitions of community response 25

S3.D. Stability of environmental canonical directions across taxonomic levels 28

S4. Phylogenetic Coherence Analysis 30

S4.A. Adjusted consenTRAIT approach 30

S4.B. Setting a Baseline for Phylogenetic Distance 33
S4.C. Pagel's $\lambda$ Approach 34

References 34

### S1. CODE AVAILABILITY

All code and pre-computed data used to generate the figures in this manuscript are deposited on GitHub (https://github.com/MilenaCW/microbiome-phylo-coherence) and archived on Zenodo (doi: 10.5281/zenodo.20586902).

### S2. DATA CURATION

#### S2.A. Selection of Datasets

##### *S2.A.1. Soil Dataset*

Soil microbial composition and environmental metadata were obtained from Delgado-Baquerizo *et al.* [1], which reports a global survey of the dominant soil bacteria. 16S ribosomal RNA gene amplicons were sequenced on the Illumina platform using paired-end chemistry. Raw sequencing files and the accompanying environmental metadata spreadsheet were downloaded from the Figshare archive associated with the original publication.

##### *S2.A.2. Ocean Dataset*

Ocean microbial composition data were obtained from the TARA Oceans expedition [2]. For each sample, metagenomically tagged 16S rRNA gene reads (miTAGs) were extracted from shotgun metagenomic data and assigned to operational taxonomic units (OTUs) at 97% sequence identity against the SILVA database, as described in Logares *et al.* [3]. The resulting OTU count table (miTAG.taxonomic.profiles.release.tsv) and the main environmental metadata (OM.CompanionTables.xlsx, Table W8) was downloaded from `https://ocean-microbiome.embl.de/companion.html`. Carbonate chemistry environmental metadata were obtained from the PANGEA data server [4].

#### S2.B. Preparing the Environmental Data

##### *S2.B.1. Soil Dataset*

The full set of candidate environmental variables available in the soil metadata comprised: pH, total organic carbon (C, %), total nitrogen (N, %), carbon-to-nitrogen ratio (C:N), total phosphorus (P, mg P $kg^{-1}$), soil texture (clay + silt, % of fine fractions), aridity index, mean diurnal temperature range, maximum and minimum temperature, precipitation seasonality, UV intensity, net primary productivity (NPP), and several ecosystem-type classifiers (continent, forest/grassland/shrubland indicators).

All Earth observation/meteorological variables (aridity, temperature, precipitation seasonality, UV, NPP) and ecosystem classifiers were excluded from the CCA, as the analysis focused on variables directly measured from soil cores. The remaining variables were filtered on the basis of high pairwise correlation, assessed by visual inspection of the correlation matrix. The C:N ratio was removed because it is a derived quantity dependent on soil C and soil N individually. The final variable set was: pH, soil C, soil N, soil P, and clay-silt fraction (Fig. S1).

Outliers were removed by applying per-variable range filters set by inspection of the marginal distributions. Samples with values exceeding the following thresholds were excluded:

$$\text{soil C} > 15\% \qquad \text{soil N} > 0.6\% \qquad \text{soil P} > 1500 \text{ mg P kg}^{-1}$$

The final soil dataset used for all analysis included $N = 217$ out of 237 original samples.

##### *S2.B.2. Ocean Dataset*

The full set of candidate environmental variables comprised core measurements from [2], depth (m), temperature (deg C), salinity (PSU), dissolved oxygen concentration (μmol/kg), phosphate concentration ($PO_4^{3-}$, μmol/L), nitrite concentration ($NO_2^-$, μmol/L), nitrite + nitrate concentration ($NO_2^- + NO_3^-$, μmol/L), silica concentration (Si, μmol/L), and carbonate chemistry parameters from the TARA Oceans carbonate dataset [4], pH, carbon dioxide concentration ($CO_2$, μmol/kg), partial pressure of carbon dioxide ($pCO_2$, μatm), fugacity of carbon dioxide ($fCO_2$, μatm), bicarbonate ion concentration ($HCO_3^-$, μmol/kg),

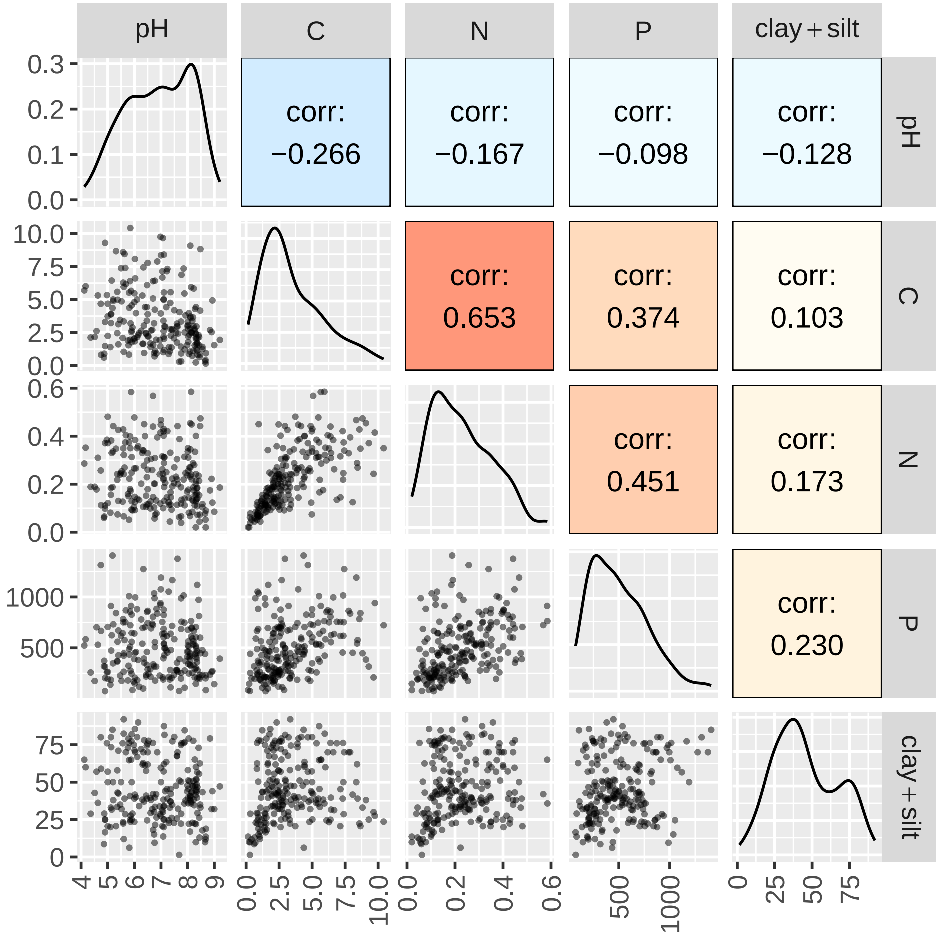


FIG. S1. **Correlogram of filtered soil environmental variables.** Pairwise relationships among five measured soil variables, pH, soil carbon (%), soil nitrogen (%), soil phosphorus (mg P/kg soil), and clay + silt texture (%), after outlier filtering (soil C $\leq$ 15%, soil N $\leq$ 0.6%, soil P $\leq$ 1,500 mg/kg) and removal of samples with missing values in any variable (final sample size: $N = 217$). Upper panels show Pearson correlation coefficients ($r$) on a colour scale from blue (negative) to red (positive). Lower panels show pairwise scatterplots. Diagonal panels show marginal density distributions for each variable.

carbonate ion concentration ($CO_3^{2-}$, $\mu$mol/kg), dissolved inorganic carbon (DIC, µmol/kg), total alkalinity (TA, µmol/kg), calcite saturation state (Ω calcite), and aragonite saturation state (Ω aragonite).

All carbonate chemistry variables were excluded because they were measured for only a small subset of samples; including them would have resulted in an unacceptable reduction in sample size (Fig. S2). Among the remaining core variables, $NO_2^- + NO_3^-$ and Si were removed

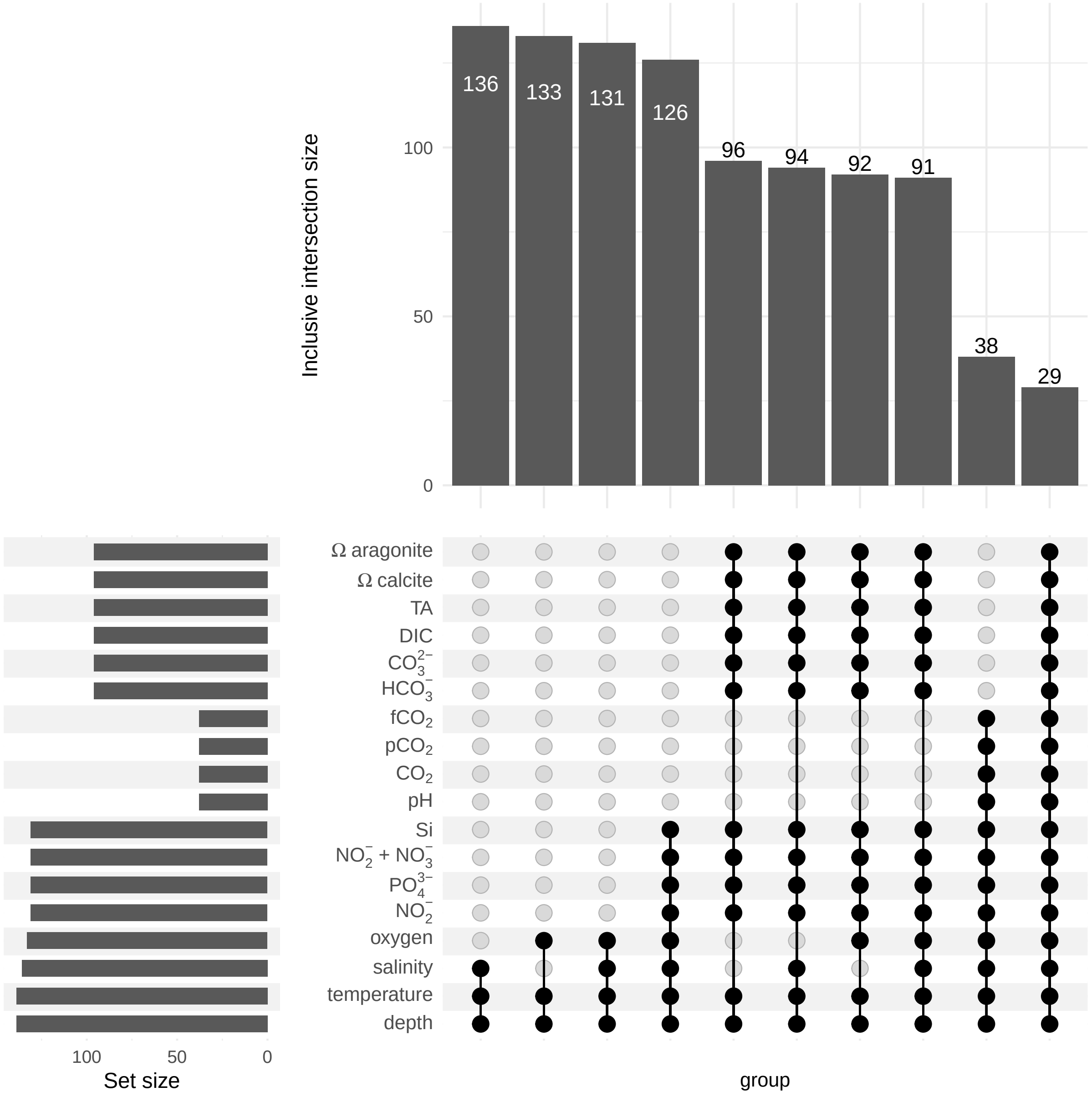


FIG. S2. **Missingness structure of the full ocean environmental variable set.** Inclusive-intersection UpSet plot showing the co-occurrence of available measurements across all candidate environmental variables: core measurements from [2] (depth, temperature, salinity, dissolved oxygen, $PO_4^{3-}$, $NO_2^-$, $NO_2^- + NO_3^-$, Si) and carbonate chemistry parameters from [4] (pH, $CO_2$, partial pressure of $CO_2$, fugacity of $CO_2$, $HCO_3^-$, $CO_3^{2-}$, dissolved inorganic carbon, total alkalinity, calcite and aragonite saturation states). Bar heights indicate the number of samples present in each intersection. Carbonate chemistry variables were available for only a small subset of samples, and were thus excluded from downstream analyses to avoid a reduction in sample size.

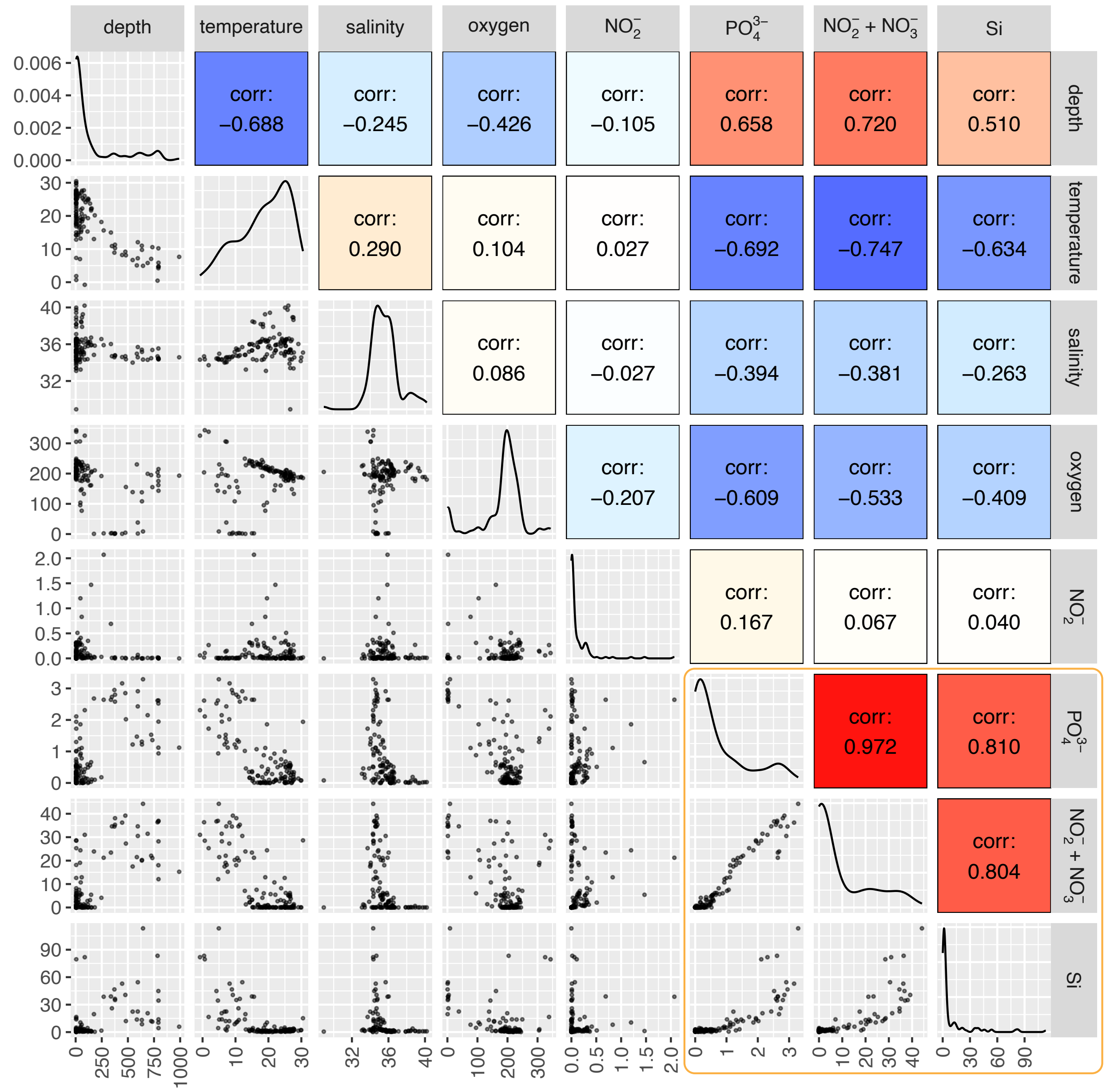


FIG. S3. **Correlogram of core ocean environmental variables.** Pairwise relationships among the eight core environmental variables from [2]: depth (m), temperature (°C), salinity (PSU), dissolved oxygen (μmol/kg), $PO_4^{3-}$ (μmol/L), $NO_2^-$ (μmol/L), $NO_2^-+NO_3^-$ (μmol/L), and Si (μmol/L). Upper panels show Pearson correlation coefficients ($r$) on a colour scale from blue (negative) to red (positive). Lower panels show pairwise scatterplots. Diagonal panels show marginal density distributions for each variable. $NO_2^-$ + $NO_3^-$ and Si were identified as highly correlated with $PO_4^{3-}$ and subsequently excluded from downstream analyses; the retained variable set comprised depth, temperature, $PO_4^{3-}$, $NO_2^-$, salinity, and dissolved oxygen.

due to high correlation with $PO_4^{3-}$, as identified by visual inspection of the correlation matrix (Fig. S3). The retained variable set was: depth, temperature, $PO_4^{3-}$, $NO_2^-$, salinity, and dissolved oxygen.

Samples with missing values in any retained variable were excluded. Additionally, samples with salinity below 32 PSU were removed as potential outliers (filters out 1 sample). The final ocean dataset used for all further analysis contains $N = 125$ samples.

#### S2.C. Preparing the Compositional Data

##### *S2.C.1. DADA2 Processing (Soil Dataset)*

Raw paired-end sequencing reads from the soil dataset were processed using the DADA2 pipeline [5]. Sequencing files were first sorted by flow cell according to the metadata in the sequence headers so that the DADA2 pipeline could be run on each flow cell independently. Reads were quality-filtered and trimmed with the following parameters: 20 bases removed from the left of each read (`trimLeft = 20`), forward reads truncated to 250 bp and reverse reads to 240 bp (`truncLen = c(250, 240)`), and reads discarded if they contained any ambiguous bases (`maxN = 0`), had more than three expected errors per read (`maxEE = c(3, 3)`), or contained a quality score below 2 (`truncQ = 2`). Expected amplicon lengths were 375–475 bp. Error rate models were learned using the DADA2 default method, followed by sample inference (denoising) and paired-end read merging. Chimeric sequences were identified and removed using the consensus method. At the end, sequence tables from each flow cell were merged into a single ASV abundance table.

##### *S2.C.2. GreenGenes2 Assignment (Both Datasets)*

*a. Preparation of sequence tables.* For the soil dataset, the merged DADA2 ASV abundance table and representative ASV sequences served as input. For the ocean dataset, the miTAG OTU count table (as reported) was used directly, bypassing the DADA2 step. Both inputs were harmonized into a standard Feature ID $\times$ sample abundance table prior to closed-reference mapping.

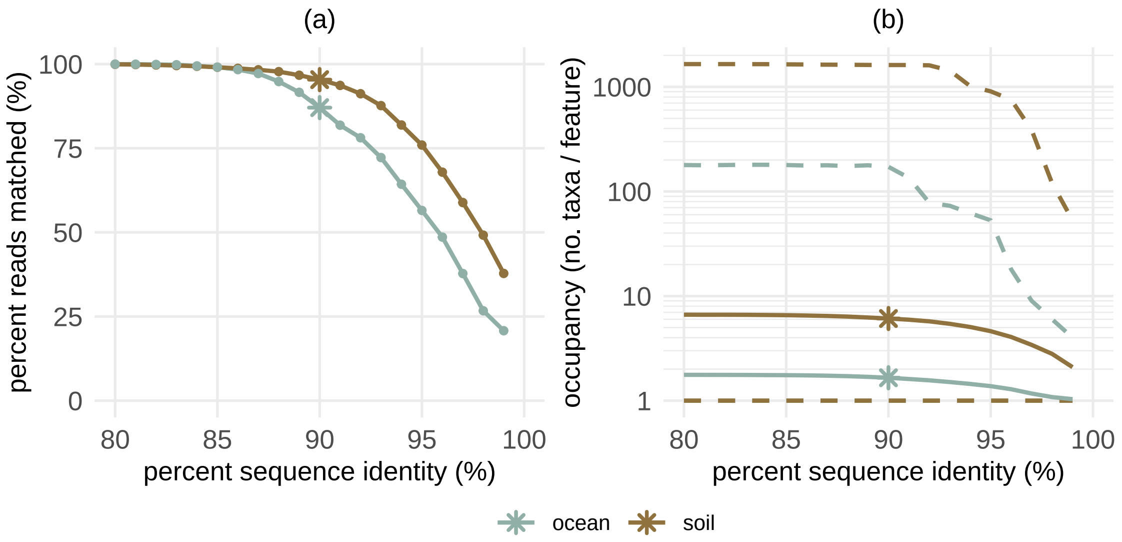


FIG. S4. **Effect of sequence identity threshold on GreenGenes2 closed-reference mapping performance for soil and ocean datasets.** Sequences were mapped to the GG2 2024.09 backbone using `vsearch` (`usearch_global` mode) across identity thresholds of 80-99%. **(a)** Percentage of reads successfully mapped as a function of identity threshold. **(b)** Occupancy, in other words the number of input ASVs (soil) or OTUs (ocean) mapped to the same GG2 backbone feature, across the same threshold range, plotted on a $\log_{10}$ scale; solid lines indicate the mean and dashed lines indicate the min/max range. Higher thresholds increase taxonomic specificity (lower occupancy) at the cost of reduced mapping success. A threshold of 90% (asterisks) was selected for both datasets as it achieved consistently high read mapping while keeping occupancy at a tractable level.

*b. Mapping and identity matching sweep.* All sequences were mapped to the GreenGenes2 (GG2) 2024.09 full-length backbone [6] using `vsearch` [7] in `usearch_global` mode (closed-reference mapping). To assess the trade-off between assignment success rate and taxonomic specificity, mapping was performed at identity thresholds ranging from 0.80 to 0.99 in 0.01 increments (Fig. S4). Higher thresholds increase the specificity of taxonomic placement (lowering occupancy or the number of ASVs/OTUs assigned to each GG2 feature) but reduce the fraction of reads successfully mapped. A threshold of 0.90 was selected for downstream analysis for both datasets as it achieved high mapping success while maintaining adequate taxonomic resolution.

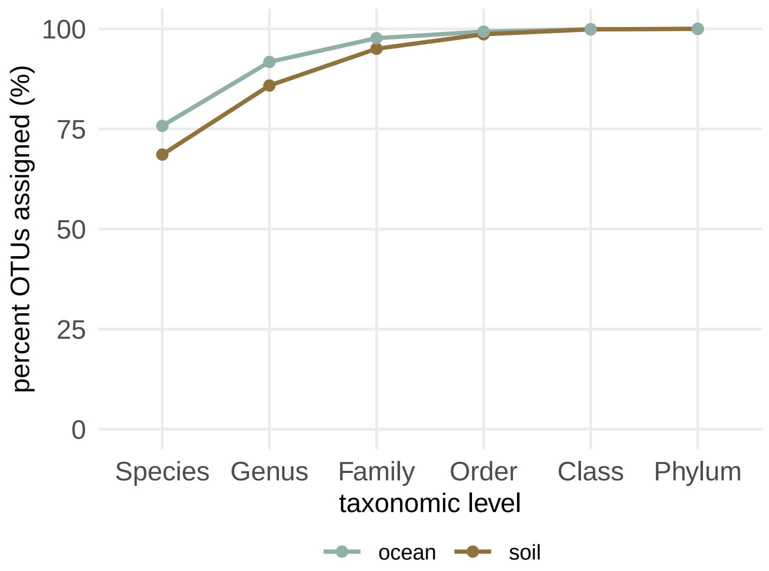


FIG. S5. **Percent of OTUs assigned taxonomy at each rank.** For each dataset, the percentage of OTUs with taxon assigned (GreenGenes2, 90% sequence identity) is shown from the finest (species) to coarsest (phylum) rank.

*c. Taxonomy and phylogeny.* The GG2 reference taxonomy (Domain through Species) was propagated to all mapped features. Note that the species-level taxonomic assignment is incomplete for a substantial fraction of OTUs (Fig. S5), as a result, species-level analyses have been excluded from all downstream cross-taxonomic comparisons. Domain-level analyses were also excluded because the classification is too coarse to be informative. The GG2 backbone phylogenetic tree was pruned to the set of mapped features. Final outputs for each dataset were: a feature count table (`seqtab.csv`), a taxonomy table (`taxonomy.csv`), and a pruned phylogenetic tree (`tree.nwk`).

*d. Structure of the GG2 tree and OTU abundances.* Two features of the pruned GG2 tree constrain what taxonomy-based null models can reveal. First, group sizes are highly skewed at every rank: the largest 10% of taxa contain 59 to 87% of OTUs, depending on rank and dataset (Fig. S6a). Second, OTU abundances are heavy-tailed: mean relative abundances span several orders of magnitude, and 2% (soil) and 6% (ocean) of OTUs account for half of all reads (Fig. S6b). Because coarse-graining sums abundances within a taxon, these properties determine how many OTUs effectively contribute to each group.

We quantify the effective contribution of OTUs by comparing the “nominal” (i.e. true) and "effective" group size. The effective group size for a taxon is defined as

$$n_{\text{effective}} = 1/\sum_i p_i^2, \tag{S1}$$

where $p_i$ is the share of the taxon's mean relative abundance carried by OTU $i$ (the inverse Simpson index). The effective size equals the nominal number of OTUs when all members are equally abundant and approaches one when a single OTU dominates. The median effective group size at genus rank is 1.3 (soil) and 1.1 (ocean), compared with a nominal size of 2, and a single OTU carries most of the abundance of a typical genus (Fig. S6c). When one OTU dominates each group, summing is close to an identity operation, so reassigning OTUs among taxa changes group-level abundances little. Randomizing OTU-to-taxon membership therefore has limited power to disrupt phylogenetic structure in this data, and we do not use it as a null model.

Note, here, and in all downstream analyses, OTUs present in fewer than two samples above a minimum read count of 10 were removed.

### S3. CANONICAL CORRELATION ANALYSIS

#### S3.A. Technical Definitions

Throughout, let $C \in \mathbb{R}^{N\times p}$ denote the compositional data matrix ($N$ samples, $p$ taxa) and $E \in \mathbb{R}^{N\times q}$ the environmental data matrix ($q$ measurements), both mean-centered. In our setting, $p \gg N$ (many more taxa than samples) while $q \ll N$ (many fewer environmental measurements than samples). The derivations in this section are adapted from [8].

##### *S3.A.1. Canonical Correlation Analysis*

The goal of CCA is to find a pair of loading vectors $\mathbf{b}_C^{(j)} \in \mathbb{R}^p$ and $\mathbf{b}_E^{(j)} \in \mathbb{R}^q$ such that the Pearson correlation between the projected data,

$$\rho^{(j)} = \text{corr}\Big(C\,\mathbf{b}_C^{(j)},\ E\,\mathbf{b}_E^{(j)}\Big), \tag{S2}$$

is maximized, subject to the constraint that successive pairs of canonical directions $\Big(\mathbf{b}_C^{(j)}, \mathbf{b}_E^{(j)}\Big)$ are orthogonal in the sense that the corresponding canonical variates are uncorrelated:

$$\text{Cov}\Big(C\,\mathbf{b}_C^{(j)},\, C\,\mathbf{b}_C^{(j')}\Big) = \text{Cov}\Big(E\,\mathbf{b}_E^{(j)},\, E\,\mathbf{b}_E^{(j')}\Big) = 0 \quad \forall\, j' < j. \tag{S3}$$

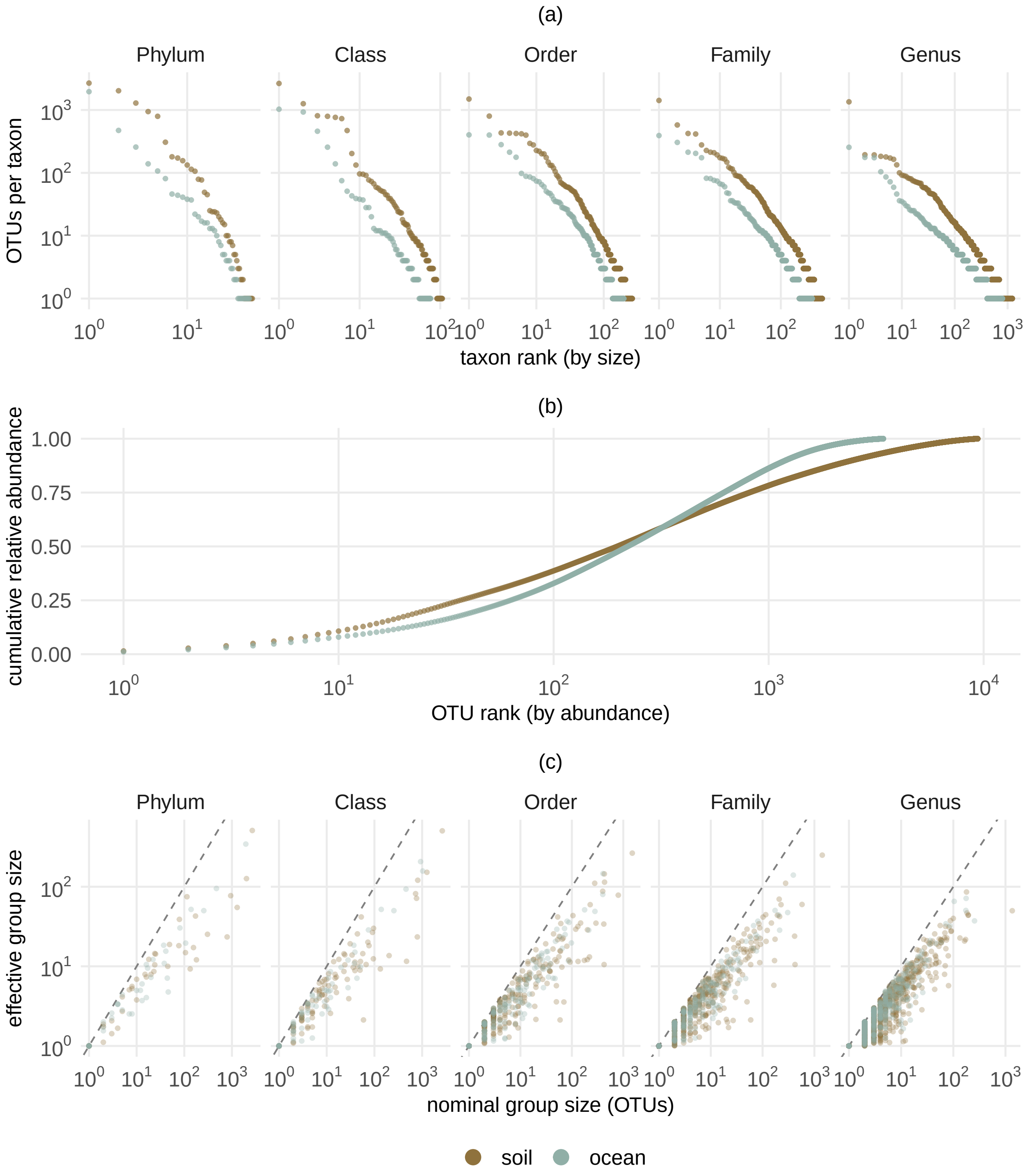


FIG. S6. **Skew in the GG2 tree and OTU abundances.** **(a)** Number of OTUs per taxon, with taxa ranked by size, at each taxonomic rank. **(b)** Cumulative mean relative abundance of OTUs ranked from most to least abundant. **(c)** Nominal group size (number of OTUs per taxon) versus effective group size (inverse Simpson index of within-taxon abundance shares), one point per taxon; the dashed line marks equality. Soil and ocean datasets at 90% sequence identity, after removal of rare OTUs.

Writing the sample covariance matrices as $\widehat{\Sigma}_{CC} = \frac{1}{N}C^T C$, $\widehat{\Sigma}_{EE} = \frac{1}{N}E^T E$, and $\widehat{\Sigma}_{CE} = \frac{1}{N}C^T E$, expanding the correlation gives

$$\left(\mathbf{b}_C^{(j)}, \mathbf{b}_E^{(j)}\right) = \underset{\mathbf{b}_C^{(j)}, \mathbf{b}_E^{(j)}}{\arg\max} \left[ \frac{\mathbf{b}_C^{(j)T} \widehat{\Sigma}_{CE}\, \mathbf{b}_E^{(j)}}{\sqrt{\mathbf{b}_C^{(j)T} \widehat{\Sigma}_{CC}\, \mathbf{b}_C^{(j)}}\, \sqrt{\mathbf{b}_E^{(j)T} \widehat{\Sigma}_{EE}\, \mathbf{b}_E^{(j)}}} \right]. \tag{S4}$$

You can then do a change of variables $\tilde{\mathbf{b}}_C^{(j)} = \widehat{\Sigma}_{CC}^{1/2}\, \mathbf{b}_C^{(j)}$ and $\tilde{\mathbf{b}}_E^{(j)} = \widehat{\Sigma}_{EE}^{1/2}\, \mathbf{b}_E^{(j)}$ to transform this into

$$\underset{\tilde{\mathbf{b}}_C^{(j)}, \tilde{\mathbf{b}}_E^{(j)}}{\arg\max} \left[ \frac{(\tilde{\mathbf{b}}_C^{(j)})^T M_{\text{CCA}}\, \tilde{\mathbf{b}}_E^{(j)}}{\|\tilde{\mathbf{b}}_C^{(j)}\|\, \|\tilde{\mathbf{b}}_E^{(j)}\|} \right], \qquad M_{\text{CCA}} = \widehat{\Sigma}_{CC}^{-1/2}\, \widehat{\Sigma}_{CE}\, \widehat{\Sigma}_{EE}^{-1/2}. \tag{S5}$$

The maximization problem can now be simply solved as an SVD problem. In other words, $\tilde{\mathbf{b}}_C^{(j)}$ and $\tilde{\mathbf{b}}_E^{(j)}$ end up as the $j$-th left and right singular vectors of $M_{\text{CCA}}$, with the corresponding singular values giving the canonical correlations $\rho^{(j)}$. The canonical directions are then recovered as $\mathbf{b}_C^{(j)} = \widehat{\Sigma}_{CC}^{-1/2} \tilde{\mathbf{b}}_C^{(j)}$ and $\mathbf{b}_E^{(j)} = \widehat{\Sigma}_{EE}^{-1/2} \tilde{\mathbf{b}}_E^{(j)}$.

#### *S3.A.2. Regularized CCA*

When $p > N$, the sample covariance matrix $\widehat{\Sigma}_{CC}$ is singular and there is risk of overfitting. One way to remedy this is by adding a diagonal regularization term (a method known as regularized CCA or RCCA [8]):

$$\widehat{\Sigma}_{CC}(\lambda_1) = \widehat{\Sigma}_{CC} + \lambda_1 I_p. \tag{S6}$$

For the environmental data, $q \ll N$ so $\widehat{\Sigma}_{EE}$ is well-conditioned; we therefore set $\lambda_2 = 0$ throughout. The RCCA canonical correlation is then

$$\rho_{\text{RCCA}}^{(j)}\left(\mathbf{b}_C^{(j)}, \mathbf{b}_E^{(j)}; \lambda_1\right) = \frac{\mathbf{b}_C^{(j)T} \widehat{\Sigma}_{CE}\, \mathbf{b}_E^{(j)}}{\sqrt{\mathbf{b}_C^{(j)T} (\widehat{\Sigma}_{CC} + \lambda_1 I)\, \mathbf{b}_C^{(j)}}\, \sqrt{\mathbf{b}_E^{(j)T} \widehat{\Sigma}_{EE}\, \mathbf{b}_E^{(j)}}}, \tag{S7}$$

and canonical directions are obtained from the SVD of

$$M_{\text{RCCA}} = \left(\widehat{\Sigma}_{CC} + \lambda_1 I\right)^{-1/2} \widehat{\Sigma}_{CE}\, \widehat{\Sigma}_{EE}^{-1/2}. \tag{S8}$$

The regularization parameter $\lambda_1$ can be interpreted as an $\ell_2$ (ridge) penalty that constrains $\|\mathbf{b}_C^{(j)}\|^2 \le t_1$ for some $t_1$ that decreases as $\lambda_1$ increases.

#### S3.B. Computational Implementation

##### *S3.B.1. Data Preprocessing*

Prior to CCA, taxa present in fewer than two samples above a minimum read count of 10 were removed. Remaining counts were converted to relative abundances (per-sample normalization). Both the composition matrix $C$ and the environmental matrix $E$ were standardized to zero mean and unit variance per feature (z-score normalization).

##### *S3.B.2. K-fold Cross-Validation and Hyperparameter Search*

The regularization parameter $\lambda_1$ was selected by $k$-fold cross-validation with $k = 15$ folds (Fig. S7a). Fold indices were assigned by random permutation of sample indices (seed = 1 for reproducibility), yielding approximately equal fold sizes. For each fold, $f$, RCCA was trained on the remaining $k - 1$ folds and evaluated on the held-out fold by computing the out-of-sample Pearson correlation between the canonical variates:

$$\rho_{\text{oos}}^{(j,f)}(\lambda_1) = \text{corr}\Big(C_{\text{test}}^{(f)}\,\mathbf{b}_C^{(j,f)}(\lambda_1),\ E_{\text{test}}^{(f)}\,\mathbf{b}_E^{(j,f)}(\lambda_1)\Big)\,. \tag{S9}$$

A grid of 21 values of $\lambda_1$ log-spaced between $10^{-5}$ and $10^5$ (soil) or between $10^{-2.5}$ and $10^5$ (ocean) was evaluated in parallel. The optimal $\lambda_1^*$ was chosen as the value maximizing the mean out-of-sample correlation of the first canonical direction ($j = 1$) across all $k$ folds:

$$\lambda_1^* = \arg\max_{\lambda_1} \frac{1}{k}\sum_{f=1}^{k} \rho_{\text{oos}}^{(1,f)}(\lambda_1) \tag{S10}$$

(Fig. S7b).

##### *S3.B.3. Loadings Extraction, Normalization, and Alignment*

At the optimal $\lambda_1^*$, RCCA was trained separately on each set of $k-1$ training folds, yielding $k$ sets of canonical directions. The resulting loading vectors $\mathbf{b}_C^{(j,f)}$ and $\mathbf{b}_E^{(j,f)}$ (for canonical direction $j$ and fold set $f$) were normalized to unit length. Because CCA is invariant to the sign of the loading vectors, sign ambiguity was resolved by aligning the environmental loadings across folds: the sign of each fold's environmental loading vector was chosen to

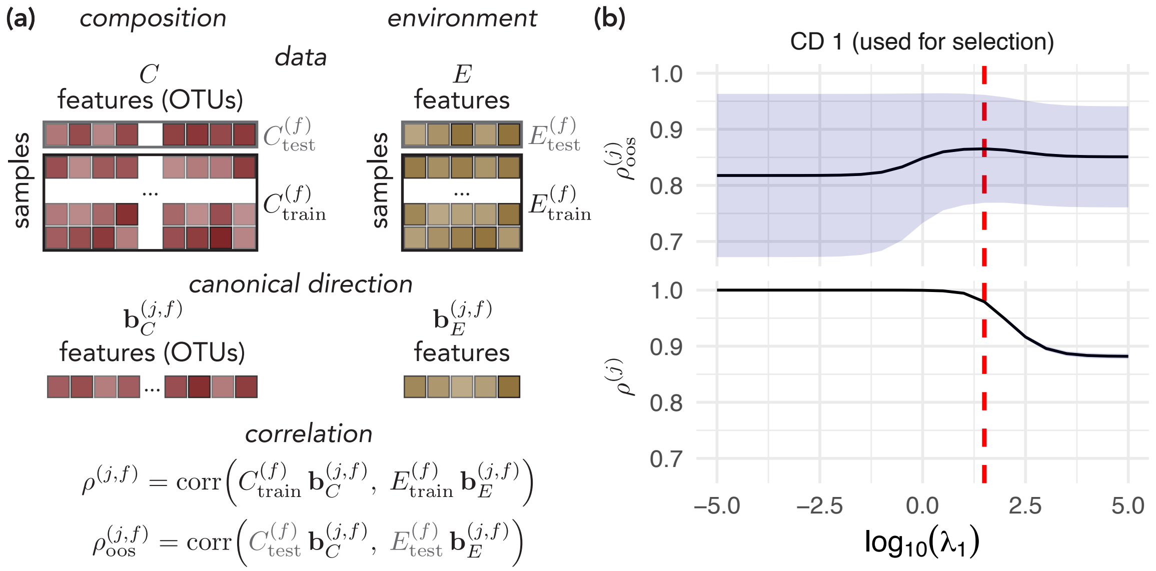


FIG. S7. **Cross-validation procedure and hyperparameter selection for RCCA. (a)** Schematic of the $k$-fold cross-validation scheme. For each fold $f$, the composition matrix $C$ and environmental matrix $E$ are partitioned into training and held-out test sets. RCCA is fit on the training data to obtain canonical directions $\mathbf{b}_C^{(j,f)}$ and $\mathbf{b}_E^{(j,f)}$, and performance is assessed by computing both the in-sample correlation $\rho^{(j,f)}$ and the out-of-sample correlation $\rho^{(j,f)}_{\text{oos}}$ on the held-out fold. **(b)** Example mean fold correlations for the first canonical direction ($j = 1$) as a function of the regularization parameter $\lambda_1$, illustrated using the soil dataset at the OTU taxonomic resolution. The upper panel shows the out-of-sample correlation $\rho^{(1)}_{\text{oos}} = \frac{1}{k}\sum_f \rho^{(1,f)}_{\text{oos}}$, with the shaded band indicating $\pm 1$ standard deviation across folds; the optimal $\lambda_1^*$ (red dashed line) is chosen as the value maximizing this mean. The lower panel shows the corresponding in-sample correlation $\rho^{(1)} = \frac{1}{k}\sum_f \rho^{(1,f)}$, which remains near unity for small $\lambda_1$ and decreases as regularization increases, consistent with the expected bias–variance trade-off.

maximize its correlation with the first fold's corresponding vector (Fig. S8), and the same sign flip was applied to the compositional loadings. Canonical directions were ranked by their mean out-of-sample correlation across folds, with $j = 1$ denoting the direction with the highest correlation.

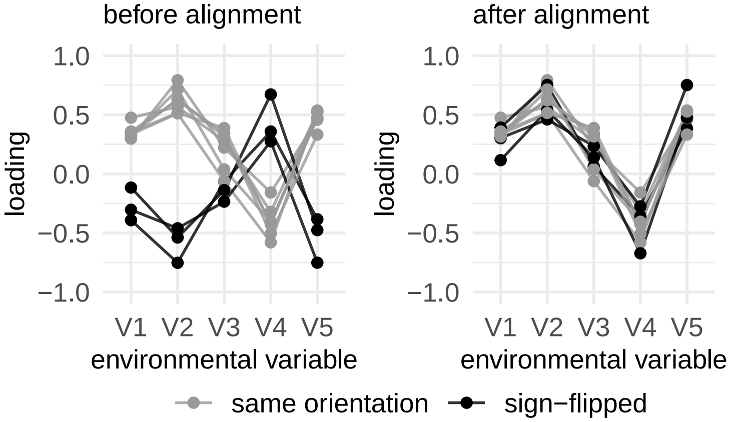


FIG. S8. **Example of sign-flip alignment of environmental loading vectors across cross-validation folds.** Approach illustrated using synthetic data. Environmental loading vectors from each cross-validation fold are plotted before (left) and after alignment (right). Each line represents one fold; folds whose loading vectors were sign-flipped to match the orientation of the first fold are shown in black, while folds that retained their original orientation are shown in grey.

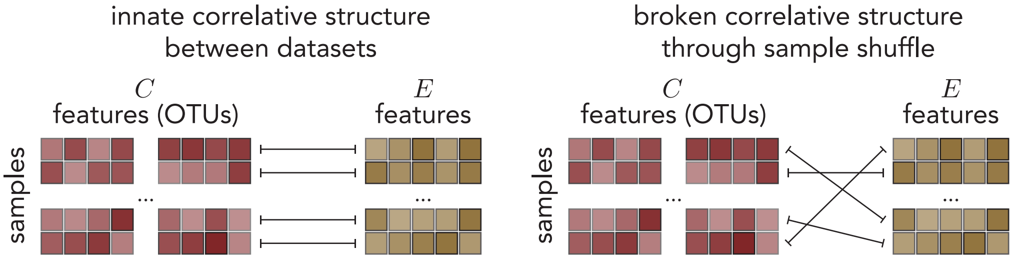


FIG. S9. **Schematic illustration of the sample-shuffle null.** For each null replicate, rows of $C$ are randomly permuted (right), breaking the sample-level pairing between composition and environment while preserving within-dataset structure.

*S3.B.4. Null Model*

To assess whether the observed CCA correlations exceed what would arise by chance, a null distribution was constructed by shuffling the rows of $C$ (the composition matrix) while leaving $E$ unchanged. This procedure preserves the internal covariance structure within each individual dataset, but destroys any correspondence between an individual sample's microbial composition and its environmental measurements.

For each of 101 independent shuffle replicates (seeds 0-100), a full $k$-fold hyperparameter

TABLE S1. **Fraction of cross-validation folds at or below the null.** Number of folds (out of $k = 15$) whose out-of-sample correlation is $\leq \bar{\rho}_{\text{null}}$, with the corresponding $p^{(j)}$ in parentheses (0.90 identity, OTU level). Asterisks denote directions for which no fold falls at or below the null ($p^{(j)} < 1/k$).

| | CD1 | CD2 | CD3 | CD4 | CD5 | CD6 |
|---|---|---|---|---|---|---|
| Soil ($\bar{\rho}_{\text{null}} = 0.18$) | $0^*$ | $0^*$ | $0^*$ | $0^*$ | 2 (0.13) | – |
| Ocean ($\bar{\rho}_{\text{null}} = 0.30$) | $0^*$ | $0^*$ | $0^*$ | 3 (0.20) | 3 (0.20) | 4 (0.27) |

search was performed on the shuffled data, and loadings and correlations were extracted at the shuffle's own optimal $\lambda_1$. Within each replicate, canonical directions are ranked by descending out-of-sample correlation, exactly as in the true fit. We compare each true canonical direction $j$ against the same reference, each null replicate's top-ranked (first) canonical direction, i.e. the strongest correlation attainable by chance in that replicate, since later null-ranked directions are, by construction, weaker and not independent estimates of chance correlation at that rank. For each replicate, the out-of-sample correlation of this first canonical direction is averaged across the $k = 15$ cross-validation folds, giving one value per replicate. The true correlations are then compared against the mean $\pm$ standard error (SE is sufficiently small that the error range is not visible) of these 101 per-replicate values (i.e. an average across fold-averages; main text Fig. 2c, SI Fig. S10).

To quantify how consistently each true canonical direction exceeds the null, we compute, for each direction $j$, the fraction of cross-validation folds whose out-of-sample correlation falls at or below the null mean defined above,

$$p^{(j)} = \frac{\#\{f : \rho^{(j)}_{\text{oos},f} \leq \bar{\rho}_{\text{null}}\}}{k} \tag{S11}$$

where $\rho^{(j)}_{\text{oos},f}$ is the out-of-sample correlation of direction $j$ on held-out fold $f$ and $\bar{\rho}_{\text{null}}$ is the mean over the 101 replicates of the fold-averaged first null canonical direction. Each fold is a disjoint subset of held-out samples, so $p^{(j)}$ can be read as the proportion of held-out subsets on which the true coupling is no better than chance. Because $p^{(j)}$ is resolved in steps of $1/k$, its smallest nonzero value is $1/15 \approx 0.067$; when no fold falls at or below the null we therefore report $p^{(j)} < 1/k$ rather than $p^{(j)} = 0$, and mark the direction with an asterisk (main text Fig. 2c). Values for all canonical directions are listed in Table S1.

##### *S3.B.5. CCA at Different Taxonomic Levels*

To assess the sensitivity of CCA results to taxonomic resolution, the analysis was repeated at multiple levels. Relative abundances of OTUs were summed within each taxonomic group (Phylum, Class, Order, Family, Genus, OTU) per sample. Given taxonomic ranks correspond to nested clades of increasing phylogenetic depth (OTU $\subset$ genus $\subset$ family $\subset \ldots \subset$ phylum), aggregating OTUs at each taxonomic level provides a coarse, discrete test as classic phylogenetic analyses that resolve a more continuous coherence by traversing the branches of the tree (e.g. consenTRAIT [9] and Pagel's $\lambda$ [10]). If closely related OTUs share similar environmental responses, the composition-environment coupling structure should survive aggregation, whereas phylogenetically incoherent responses should average out and distort the coupling.

CCA was run independently at each level using the same pipeline and hyperparameters described above (out-of-sample correlation performance across taxonomic levels shown in Fig. S10). The null distribution shown for each taxonomic level in Fig. S10 is the same sample-shuffle null described above (Sec. S3.B.4), applied independently to that level's own aggregated composition matrix.

#### S3.C. Relation to other methods

##### *S3.C.1. PCA vs. CCA decomposition of community composition*

The most common approach to examine the relationship between community composition and environmental variation in global survey datasets is to start with a linear decomposition of the relative abundance matrix via principle component analysis (PCA) and then ask how each environmental variable correlates with each of the leading principle components (e.g. [2, 11, 12]). This method allows you to conclude whether certain environmental variables are more or less correlated with axes of high variation in composition, however it is limited in that it does not utilize all of the mutual information that is contained across both the compositional and environmental data. By using solely the compositional data to decompose community composition, you are assuming that the variation in composition due to the environment overpowers any other sources of variation (e.g. community dynamics, interactions, and noise) [13]. Secondarily, this method also assumes that composition is

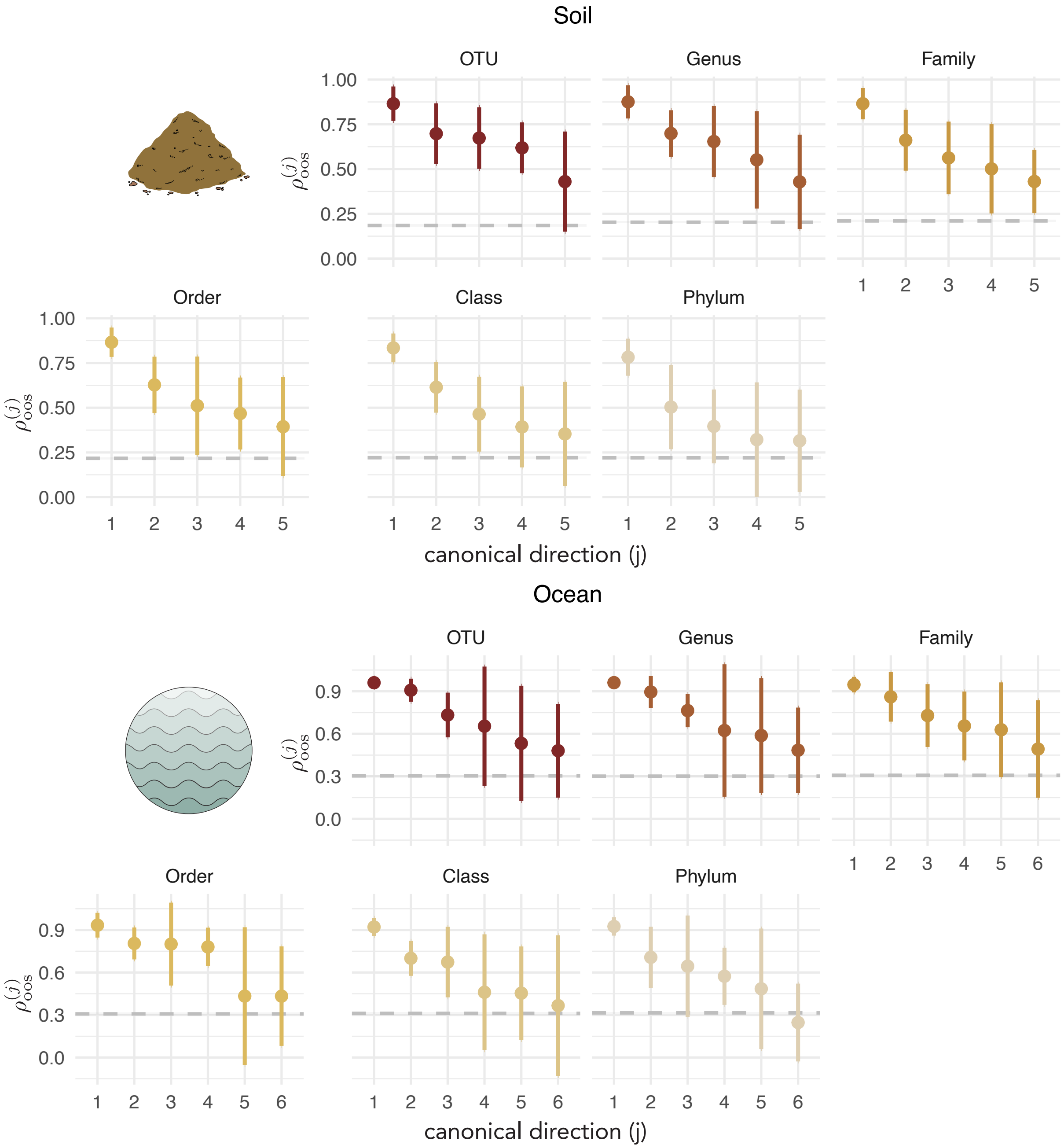


FIG. S10. **Out-of-sample CCA correlation across taxonomic levels for soil and ocean datasets.** Out-of-sample canonical correlation ($\rho^{(j)}_{\mathrm{oos}}$, mean $\pm$ SD across cross-validation folds) is shown for each canonical direction $j$, evaluated separately at six taxonomic levels (OTU, Genus, Family, Order, Class, Phylum), for the soil dataset (top) and ocean dataset (bottom). The dashed gray line indicates the null distribution threshold (mean $\pm$ SE across sample-shuffled permutations, using each replicate's first canonical direction as the reference, as in Fig. 2c).

varying with pure axes of environmental change and ignores/disallows correlations that may fall along multivariate axes of environmental variation.

By using CCA, our approach specifically looks for variation in the composition coupled to environmental variation through maximizing the correlation (see details on the method above in SI Sec. S3.A). By definition, CCA uses all of the information at hand to decompose how microbial composition and environment *co-vary*. It also allows for the possibility that the processes driving compositional change (trait or otherwise) are multivariate. This is relevant given how interconnected and correlated the different environmental measurements are with one another both mechanistically and statistically. Among the five retained soil variables, the maximum absolute pairwise correlation is $r_{\max} = 0.65$ (between total organic carbon and total nitrogen), and among the six retained ocean variations, $r_{\max} = -0.688$ (between depth and temperature) (SI Figs. S1 and S3).

To directly compare how the results of our CCA approach could diverge from a classic PCA approach, we characterize how the canonical directions in compositional space relate to the dominant axes of compositional variation. We specifically compared the direction and explained variance of each canonical direction (CD) against the principal components (PCs) of the composition matrix.

*a. Angle with PC1.* For each cross-validation fold $f$ and each canonical direction $j$, the CCA abundance loading vector $\mathbf{b}_C^{(j,f)}$ was compared against the first principal component, $\mathrm{PC1}^{(f)}$, as computed from the training composition matrix $C_{\mathrm{train}}^{(f)}$ (already mean centered and variance normalized) using PCA (`prcomp`). The angle between each CD and PC1 is

$$\theta^{(j,f)} = \arccos\Big(\big|\langle \hat{\mathbf{b}}_C^{(j,f)}, \mathrm{PC1}^{(f)} \rangle\big|\Big) ,$$

where the absolute value accounts for the sign indeterminacy of eigenvectors.

To construct a null distribution for $\theta^{(j,f)}$, the abundances of each sample in $C_{\mathrm{train}}^{(f)}$ were independently permuted across taxa for 100 replicates, and PCA was recomputed on each permuted matrix. This preserves each sample's marginal distribution of relative abundances across the full taxon set while destroying inter-taxon co-abundance structure. PC1 was computed for each shuffled matrix, and the angle between each CD and the resulting null PC1 was recorded. The null thus captures what angle is expected when PC1 reflects only the per-sample diversity profile, with no consistent cross-sample covariance structure.

The angle between CD1 and PC1 (color) falls substantially below the null distribution

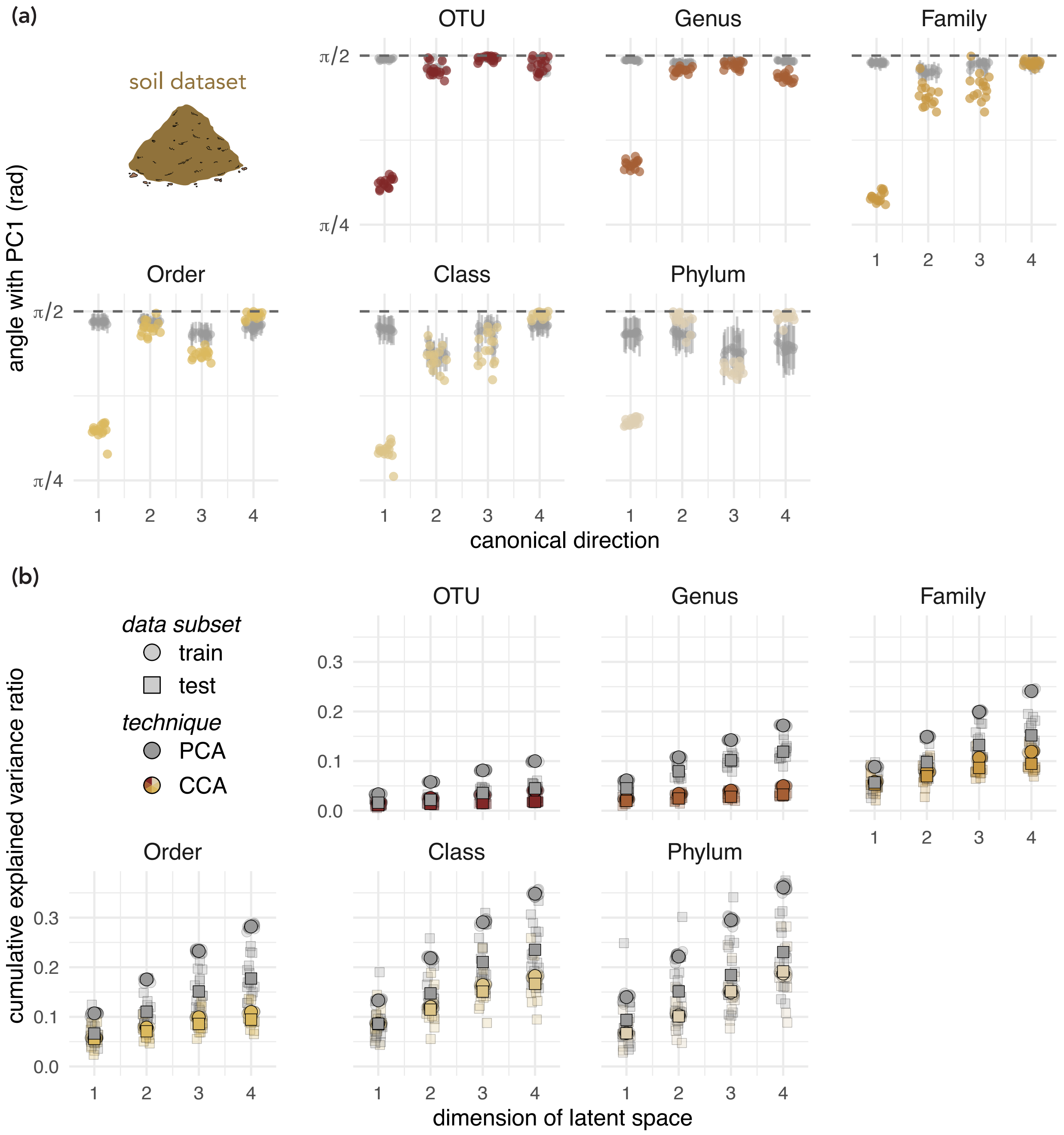


FIG. S11. **Comparison of CCA against PCA for the soil dataset.** Results are shown for each of the significant canonical directions, evaluated across cross-validation folds and six taxonomic levels (OTU through Phylum; color scale from dark red to tan). **(a)** Angle (radians) between each canonical direction (CD) and first principal component (PC1) of the training composition matrix. Colored points show the true angle per fold; grey point-ranges show the mean ± SD of the null distribution (100 replicates in which entries in each sample are randomly permuted before computing PC1).

FIG. S11. *Caption continued.* The dashed line marks $\pi/2$ (vectors are orthogonal). CD1 falls substantially below the null; subsequent CDs do not. **(b)** Cumulative explained variance ratio (EVR) as a function of the number of dimensions retained. Circles denote training-set EVR; squares denote test-set EVR. Grey symbols show PCA; colored symbols show CCA (orthogonalized via QR). Faint (low-opacity) points show individual folds; solid (high-opacity) points show mean across folds. PCA achieves higher in-sample EVR but exhibits a large train–test gap; CCA incurs a modest EVR cost while remaining stable across splits.

(grey) for both datasets, indicating that the first canonical direction is considerably more aligned with the dominant axis of compositional variance than expected by chance. Subsequent canonical directions have angles that are indistinguishable from the null, consistent with their role in capturing variation orthogonal to the leading axis (Fig. S11a and Fig. S12a). This tells us that, in the case of the soil and ocean datasets, CCA and PCA pull out the same primary axis of variation.

*b. Cumulative explained variance ratio.* The explained variance ratio (EVR) at dimension $d$ is the fraction of the total sum of squares of the composition matrix, $C$, captured by the first $d$ directions:

$$\mathrm{EVR}(d) = \frac{\sum_{j'=1}^{d} \|C\,\mathbf{q}_{j'}\|^2}{\|C\|_F^2},$$

where $\mathbf{q}_{j'}$ is the $j'$-th column of the orthonormal basis $Q$ under evaluation. For PCA, $Q^{(f)}_{\mathrm{PCA}}$ is the orthonormal basis matrix from `prcomp` fitted on $C^{(f)}_{\mathrm{train}}$. For CCA, the abundance loading matrix $B^{(f)}_C$ (columns: loading vectors, $\mathbf{b}^{(j,f)}_C$, for each CD $j$) is orthonormalized via QR decomposition to obtain $Q^{(f)}_{\mathrm{CCA}}$, placing CCA and PCA on a comparable footing. EVR is evaluated on both $C^{(f)}_{\mathrm{train}}$ (in-sample) and $C^{(f)}_{\mathrm{test}}$ (out-of-sample).

PCA achieves higher EVR on the training set than CCA, as expected given the PCA decomposition is the optimal orthonormal basis for EVR on the training data (both evaluated within the same fold split). However, PCA shows a consistently large gap between training and test EVR across folds and taxonomic levels, indicating that the leading PCs overfit to idiosyncratic variance in the training samples. This tells us that while CCA directions pay a modest cost in training EVR, they are more stable between train and test folds (Fig. S11b and Fig. S12b). Coupling the decomposition to the environment thus seemingly acts as an implicit regularizer, preferentially retaining variance axes that generalize across samples.

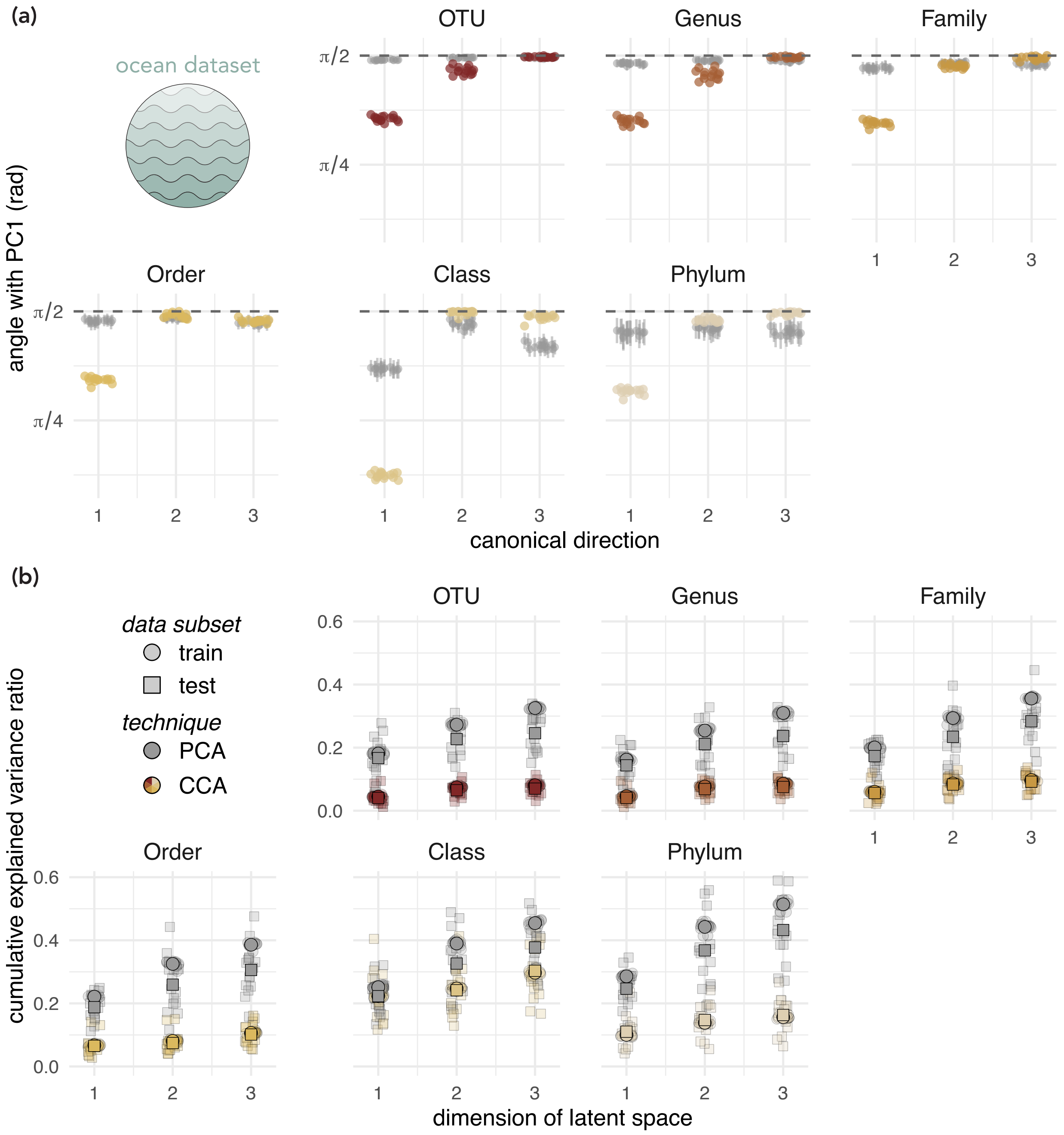


FIG. S12. **Comparison of CCA against PCA for the ocean dataset.** As in Fig. S11, for the TARA Oceans dataset. **(a)** Angle between each canonical direction (CD) and PC1, with null distribution from within-sample permutation of the training composition matrix. **(b)** Cumulative EVR for CCA (colored) and PCA (grey) evaluated on training (circles) and test (squares) folds.

*S3.C.2. Correlation-based vs. canonical weight definitions of community response*

Related to PCA decomposition and subsequent correlation with environmental variables is another method to understand which environmental directions are most relevant to community composition and phylogeny. In this approach, one computes the correlation between the abundances of each OTU across all samples and an environmental axis, then tests the phylogenetic coherence of the resulting vector of correlations.

In its standard form, the environmental axis is a single measured environmental variable, $\mathbf{e}_k$ (e.g. pH), and each OTU's response trait is $\mathrm{corr}(\mathbf{c}_i, \mathbf{e}_k)$ (see, e.g., [14]). We call this the *single-variable correlation* trait. The same calculation can be performed against any axis in environmental space. A natural choice is the $j$th environmental canonical variate from CCA, $V^{(j)} = E\mathbf{b}_E^{(j)}$, i.e. the projection of the samples onto the $j$th environmental canonical direction. The correlation between a variable in one set and a canonical variate of the other set is known as a *cross-loading*, so we refer to $\mathrm{corr}(\mathbf{c}_i, V^{(j)})$ as the *cross-loading* trait. The cross-loading trait retains the correlation-based definition of response but uses the same environmental axes as our method, making it the most direct point of comparison (Table S2).

The cross-loading trait differs from our approach only in how one defines the "response" of each OTU; the single-variable correlation trait additionally differs in its choice of environmental axis. In this paper we use the canonical weights in compositional space, $\mathbf{b}_C^{(j)}$, as our proxy for community response. Each weight reflects that OTU's contribution to the compositional signal that tracks the environmental axis $V^{(j)}$, after accounting for the contributions of all other OTUs to a degree set by the regularization strength $\lambda_1$. All three traits are alternative proxies for the same underlying quantity. Note that in all three cases the "response" is correlative and statistical rather than causal or derived from an explicit dynamical model.

*a. Limitation of the single-variable correlation trait.* The single-variable correlation trait shares the central limitation of the PCA approach as described in Sec. S3.C.1 above; it is inherently univariate, testing one environmental measurement at a time, and thus cannot capture a "response" to a jointly varying combination of correlated environmental drivers. The cross-loading trait avoids this limitation, since $V^{(j)}$ is itself a multivariate composite defined by CCA. The remaining question is how the cross-loading and canonical weight traits

TABLE S2. Three ways to define the “response” of OTU $i$ to an environmental axis. The response vector $(r_1, \dots, r_p)$ is the trait whose phylogenetic coherence is tested. $\mathbf{e}_k$ denotes the vector across samples of the $k$th measured environmental variable. $\mathbf{c}_i$ denotes the relative abundance vector across samples of the $i$th OTU. $E$ denotes the full matrix of measured environmental variables. $\mathbf{b}_E^{(j)}$ denotes the $j$th canonical direction in environmental space, and $V^{(j)}$ the corresponding canonical variate. $\mathbf{b}_C^{(j)}$ denotes the $j$th canonical direction in compositional space.

| Trait | Environmental axis | Response of OTU $i$ | Multivariate in environment | Accounts for OTU covariance |
|---|---|---|---|---|
| single-variable correlation | $\mathbf{e}_k$ | $r_i = \mathrm{corr}(\mathbf{c}_i, \mathbf{e}_k)$ | No | No |
| cross-loading | $V^{(j)} = E\mathbf{b}_E^{(j)}$ | $r_i = \mathrm{corr}(\mathbf{c}_i, V^{(j)})$ | Yes | No |
| canonical weight (this paper) | $V^{(j)} = E\mathbf{b}_E^{(j)}$ | $r_i = b_{C,i}^{(j)}$ | Yes | Yes |

differ, which we address next.

*b. Relationship between cross-loadings and canonical weights.* The cross-loading trait, $\mathrm{corr}(\mathbf{c}_i, V^{(j)})$, and the canonical weight trait, $b_{C,i}^{(j)}$, are connected by a single, regularization-dependent linear transformation. (The single-variable correlation trait uses a different environmental axis and is not covered by this relationship.)

Because the columns of $C$ and $E$ are each standardized to unit variance (Sec. S3.B), and because the environmental canonical variate $V^{(j)}$ has unit variance by construction in the RCCA formulation of Sec. S3.A (regularization is applied only on the compositional side), the correlation reduces to its numerator, the covariance:

$$\mathrm{corr}\big(\mathbf{c}_i, V^{(j)}\big) = \widehat{\mathrm{Cov}}\big(\mathbf{c}_i, V^{(j)}\big)\,. \tag{S12}$$

Collecting these covariances for all OTUs and expanding directly from the sample data,

$$\widehat{\mathrm{Cov}}\big(C, V^{(j)}\big) = \frac{1}{N} C^T E\, \mathbf{b}_E^{(j)} = \widehat{\Sigma}_{CE}\mathbf{b}_E^{(j)}. \tag{S13}$$

Recall that $(\widehat{\Sigma}_{CC}+\lambda_1 I)^{1/2}\mathbf{b}_C^{(j)}$ and $\widehat{\Sigma}_{EE}^{1/2}\mathbf{b}_E^{(j)}$ are the left and right singular vectors of $M_{\mathrm{RCCA}} = (\widehat{\Sigma}_{CC} + \lambda_1 I)^{-1/2}\widehat{\Sigma}_{CE}\widehat{\Sigma}_{EE}^{-1/2}$, with singular value $\rho_{\mathrm{RCCA}}^{(j)}$, the in-sample canonical correlation obtained from fitting RCCA on the training data. In other words,

$$M_{\mathrm{RCCA}}\,\widehat{\Sigma}_{EE}^{1/2}\mathbf{b}_E^{(j)} = \rho_{\mathrm{RCCA}}^{(j)}\,(\widehat{\Sigma}_{CC} + \lambda_1 I)^{1/2}\mathbf{b}_C^{(j)}. \tag{S14}$$

Rewriting $\widehat{\Sigma}_{CE}$ in terms of $M_{\mathrm{RCCA}}$ and applying Eq. (S14) allows us to concisely relate the cross-loading to the canonical weights and vice versa,

$$\operatorname{corr}\big(\mathbf{c}_i, V^{(j)}\big) = \rho_{\mathrm{RCCA}}^{(j)}\Big[\widehat{\Sigma}_{CC}\,\mathbf{b}_C^{(j)}\Big]_i + \rho_{\mathrm{RCCA}}^{(j)}\,\lambda_1\,b_{C,i}^{(j)}, \tag{S15}$$

$$\mathbf{b}_C^{(j)} = \frac{1}{\rho_{\mathrm{RCCA}}^{(j)}}\Big(\widehat{\Sigma}_{CC} + \lambda_1 I\Big)^{-1}\widehat{\operatorname{Cov}}\big(C, V^{(j)}\big)\,. \tag{S16}$$

Equation (S15) shows that the cross-loading trait is the canonical weight vector passed through the taxon–taxon correlation matrix $\widehat{\Sigma}_{CC}$, plus a copy of the weights scaled by $\lambda_1$. Equation (S16) gives the complementary reading: up to the constant $1/\rho_{\mathrm{RCCA}}^{(j)}$, the canonical weights are the ridge-regression coefficients obtained by regressing $V^{(j)}$ on all OTUs jointly, whereas each cross-loading is the coefficient obtained by regressing $V^{(j)}$ on that OTU alone (with unit variances, this simple regression coefficient equals the correlation). The canonical weights are therefore partial associations, analogous to coefficients in multiple regression, while the cross-loadings are marginal associations.

The regularization constant $\lambda_1$ controls how far apart the two traits are. As $\lambda_1 \to 0$, the weights approach ordinary multiple-regression coefficients and the cross-loadings differ from them by the full action of $\widehat{\Sigma}_{CC}$. Two taxa that co-vary strongly in abundance for reasons other than a shared response to $V^{(j)}$ (shared physiology, cross-feeding, correlated relative abundance inherited along the phylogeny, or a shared response to a different canonical axis) then have their cross-loadings pulled toward one another, regardless of whether their canonical weights agree. As $\lambda_1 \to \infty$, the covariance adjustment vanishes and the cross-loading trait becomes proportional to the canonical weight trait, with positive constant $\rho_{\mathrm{RCCA}}^{(j)}\lambda_1$. Since consenTRAIT depends only on the sign of the trait and Pagel's $\lambda$ is invariant to rescaling by a positive constant, the two traits then yield identical phylogenetic coherence. Note that $V^{(j)}$ is itself refit at each value of $\lambda_1$, so this comparison is always between the two traits computed against the same fitted canonical variate.

*c. Practical trade-offs.* Neither trait is free of the taxon–taxon covariance structure. It enters the cross-loadings multiplicatively and the canonical weights through an inverse. As a result, the two traits are susceptible to opposite errors. The cross-loading trait can overstate the response of taxa whose association with $V^{(j)}$ is inherited through co-abundance with responding taxa. The canonical weight trait can understate, or misattribute among taxa, the response of taxa that are largely redundant with one another (e.g. near-identical abundance profiles), since credit for their shared contribution may be split unevenly. Taxa that respond to the same environmental driver but also carry independent variation (the result of "non-environmental" processes) are not penalized in this way. It is redundancy, not shared environmental response, that the canonical weights discount, and $\lambda_1$ sets how strongly.

Both traits recognize environmental response only along measured variables. Co-abundance driven by an unmeasured environmental factor is treated like any other co-abundance, unless that factor is correlated with the variables in $E$, in which case part of its effect is absorbed into $V^{(j)}$ and attributed to the measured gradient.

The cross-loading trait is simpler to compute and more intuitive. The canonical weight trait is the more natural quantity if the question of interest is which taxa contribute to the compositional change captured by each learned canonical axis $V^{(j)}$, after accounting for their co-abundance with other taxa. Which trait yields higher phylogenetic coherence is an empirical question that depends on whether phylogenetically related taxa tend to co-vary in abundance for reasons unrelated to the environmental axis under consideration.

#### S3.D. Stability of environmental canonical directions across taxonomic levels

As described in the main text, if microbial responses are phylogenetically coherent, then the environment–composition coupling should remain stable when abundances are coarse-grained to higher taxonomic levels, since closely related OTUs are aggregated together (main text Fig. 3a). This means that the *environmental* canonical directions $\mathbf{b}_E^{(j)}$ should be stable across taxonomic resolutions: if the same biological signal is recovered at every level, CCA should identify the same dominant environmental gradients regardless of whether abundances are resolved at the OTU or phylum level. Here we quantify this cross-resolution stability of the environmental loading vectors to complement the qualitative argument in the main text (main text Fig. 3b).

To quantify the coherence of environmental canonical directions across taxonomic resolutions, we compared the $j$-th environmental loading vector $\mathbf{b}_E^{(j)}$ at each pair of taxonomic levels $(\alpha, \beta)$ using the angle

$$\theta_{\alpha\beta}^{(j)} = \arccos\left(\frac{\left|\mathbf{b}_{E,\alpha}^{(j)} \cdot \mathbf{b}_{E,\beta}^{(j)}\right|}{\|\mathbf{b}_{E,\alpha}^{(j)}\| \, \|\mathbf{b}_{E,\beta}^{(j)}\|}\right), \tag{S17}$$

where the absolute value accounts for the sign indeterminacy of canonical directions ($\theta = 0$ indicates identical loading directions; $\theta = \pi/2$ indicates orthogonality). Angles were averaged across cross-validation folds for each pair ($\bar{\theta}_{\alpha\beta}^{(j)}$), and the mean over all $\binom{6}{2} = 15$ pairs of the six taxonomic levels (OTU, genus, family, order, class, phylum) provided an overall coherence summary $\bar{\Theta}^{(j)}$ for canonical direction $j$.

To construct a null distribution, we repeated CCA 101 times on each coarsened abundance matrix after randomly permuting the sample rows, thereby breaking the microbiome–environment association while preserving the structure of the environmental predictor matrix. The null CCA was run independently at each taxonomic level, so any cross-coarsening similarity in the null reflects only estimation noise and shared environmental variable structure, not biological signal. The resulting null distributions for $\bar{\Theta}^{(j)}$ (Fig. S13) and $\bar{\theta}_{\alpha\beta}^{(j)}$ (Fig. S14) establish the angles expected in the absence of phylogenetically coherent responses.

For both datasets, the first canonical direction is highly coherent: the observed $\bar{\Theta}^{(1)}$ is far below the null distribution. Higher canonical directions show increasing overlap with the null distribution, indicating they are progressively incoherent across taxonomic resolutions; this trend is consistent with the main text's claim that secondary canonical directions are incoherent (main text Figs. 3, 4).

At the level of individual pairs (Fig. S14), nearly all 15 pairs lie below the null 95% confidence interval for the first canonical direction, confirming that this dominant environmental gradient is coherent across all pairwise combinations of taxonomic resolution. For higher canonical directions, an increasing fraction of pairs falls within the null interval; those that remain below it tend to involve adjacent or near-adjacent taxonomic levels (e.g. class–phylum), suggesting that some information about secondary gradients is retained when coarse-graining changes resolution only modestly.

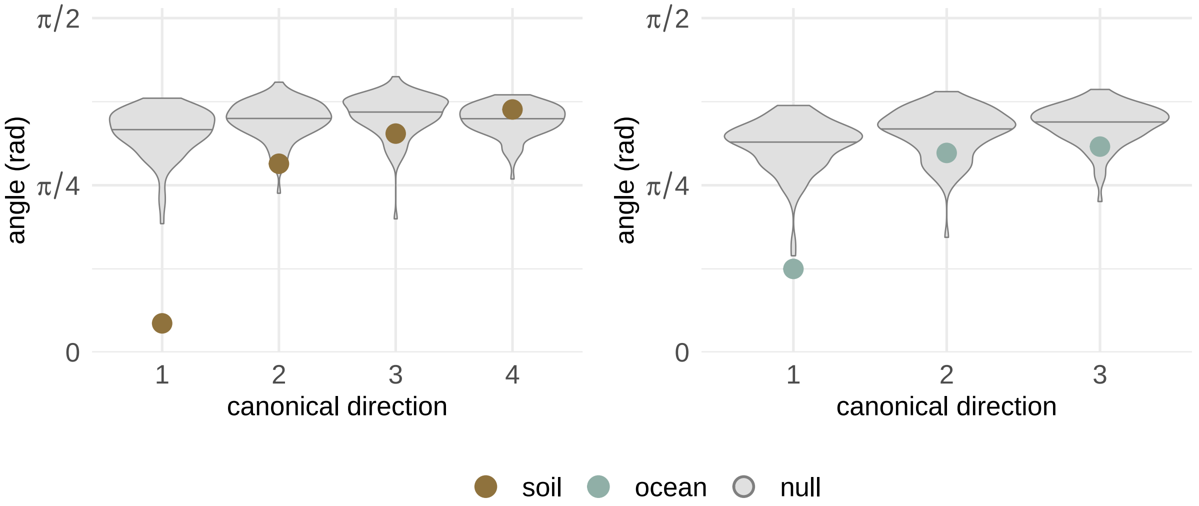


FIG. S13. **Stability of environmental canonical directions across taxonomic levels.** Mean angle $\bar{\Theta}^{(j)}$ between all pairs of taxonomic levels for each canonical direction $j$ (soil, left; ocean, right). Grey violins show the distribution of null mean angles across 101 row-permuted CCA runs; line marks the median. Colored points show the observed $\bar{\Theta}^{(j)}$ (brown: soil; teal: ocean). An angle of 0 indicates identical loading vectors; $\pi/2$ indicates orthogonal vectors.

### S4. PHYLOGENETIC COHERENCE ANALYSIS

Phylogenetic coherence analyses were carried out using OTU-level CCA results (perc-identity threshold 0.90). The GG2 backbone phylogenetic tree, pruned to OTUs present in the composition matrix, was used throughout.

#### S4.A. Adjusted consenTRAIT approach

We adapted the consenTRAIT framework [9] to quantify the phylogenetic depth of OTU clades with coherent CCA abundance-loading signs. The original consenTRAIT algorithm was designed for binary presence/absence traits: it identifies clades in which at least a specified fraction of tips share the focal trait and estimates the mean genetic depth, $\tau_D$, of those coherent clades. For signed CCA loadings, applying the original binary procedure would require two separate analyses, one treating positive loadings as the focal trait and one treating negative loadings as the focal trait. Because the sign of a CCA axis is arbitrary,

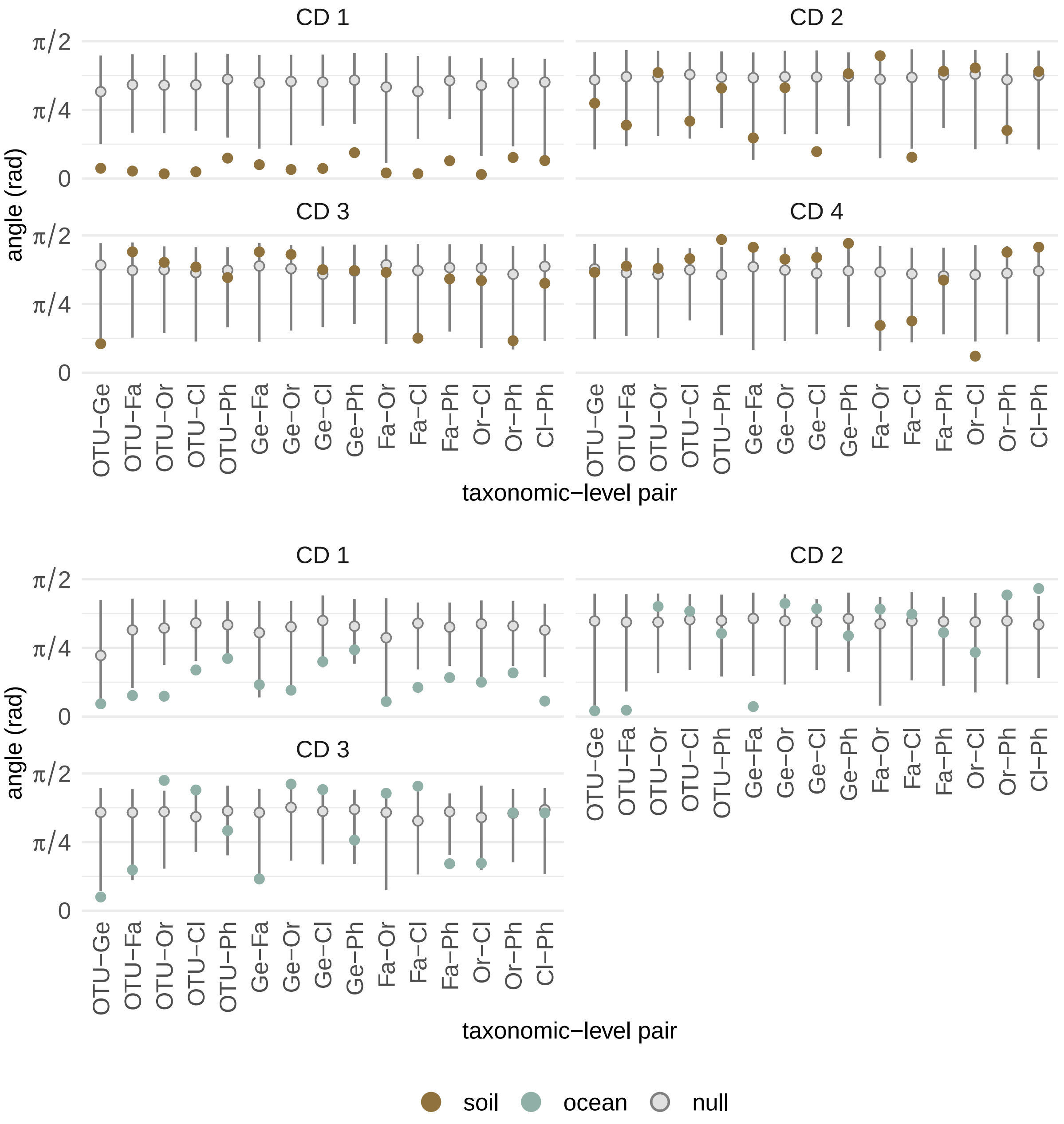


FIG. S14. **Pairwise cross-coarsening angles by canonical direction.** Each panel shows one canonical direction. The x-axis lists all 15 pairs of taxonomic levels (OTU: operational taxonomic unit; Ge: genus; Fa: family; Or: order; Cl: class; Ph: phylum). Grey points and bars show the null median and 95% confidence interval (2.5th–97.5th percentile) across 101 row-permuted CCA runs. Colored points show the observed mean angle $\bar{\theta}^{(j)}_{\alpha\beta}$ averaged across cross-validation folds (brown: soil; teal: ocean).

we instead implemented a signed version, `consentrait_signed`, that treats positive and negative loading clades symmetrically in a single analysis.

For each canonical direction $j$ and cross-validation fold $f$, we converted each nonzero OTU abundance loading to a signed trait,

$$z_i^{(j,f)} = \mathrm{sign}\Big(b_{C,i}^{(j,f)}\Big) \in \{-1, +1\},$$

where $\mathbf{b}_C^{(j,f)}$ is the CCA abundance-loading vector and $b_{C,i}^{(j,f)}$ is the loading of OTU $i$. For a candidate subtree $u$ with $n_u$ descendant tips, we counted the number of positive and negative tips, $n_+(u)$ and $n_-(u)$. The subtree was declared sign-coherent when

$$\frac{\max\{n_+(u), n_-(u)\}}{n_u} \geq \phi_{\text{consensus}} \ ,$$

where $\phi_{\text{consensus}} = 0.90$ is the consensus threshold. Equivalently, because $z_i^{(j,f)} \in \{-1, +1\}$, the criterion can be written as

$$\frac{\left|\sum_{i \in u} z_i^{(j,f)}\right|}{n_u} \geq 2\phi_{\text{consensus}} - 1.$$

This form makes the sign symmetry explicit: reversing the sign of every loading, $\mathbf{b}_C^{(j,f)} \mapsto -\mathbf{b}_C^{(j,f)}$, exchanges positive and negative tips but leaves the set of coherent clades, and therefore $\tau_D$, unchanged.

The algorithm first performs a post-order traversal of the tree to compute, for every subtree, the number of descendant tips, the signed sum of descendant trait values, and the summed root-to-tip distances of descendant tips. It then performs a pre-order traversal to identify the largest non-overlapping coherent subtrees: once a coherent subtree is selected, its descendants are not evaluated as separate coherent clades. Tips not contained within any coherent subtree are treated as singletons, following the original consenTRAIT treatment of isolated trait occurrences.

For coherent clade $k$, let $n_k$ be its number of descendant tips, let $r_k$ be its MRCA node, and let

$$d_k = \frac{1}{n_k} \sum_{i \in k} \mathrm{dist}(i, r_k)$$

be the mean distance from the clade MRCA to its descendant tips. For a singleton tip $i$, let

$\ell_i$ be its terminal branch length. The signed, size-weighted mean genetic depth is then

$$\tau_D = \frac{\sum_{k\in\mathcal{K}} n_k d_k + \alpha \sum_{i\in\mathcal{S}} \ell_i}{\sum_{k\in\mathcal{K}} n_k + |\mathcal{S}|},$$

where $\mathcal{K}$ is the set of coherent clades, $\mathcal{S}$ is the set of singleton tips, and $\alpha = 0.5$. The singleton contribution therefore follows the original consenTRAIT convention of assigning singleton entries half the terminal depth. The clade contribution, however, differs from the original implementation by weighting each coherent clade by its number of tips rather than giving each clade equal weight.

To assess significance, we generated a null distribution by randomly permuting the signed trait values across the tips of the pruned tree. This preserves the observed number of positive and negative loadings for each canonical direction and fold while breaking any association between loading sign and phylogenetic position.

Across the significant canonical directions in both datasets, the observed signed, size-weighted $\tau_D$ values were larger than the shuffled null expectation (main-text Figs. 3d, 4c). Thus, OTUs with the same sign of response along a CCA axis are not randomly distributed across the phylogeny; rather, same-sign loadings are shared within deeper phylogenetic clades than expected after permuting loading signs across tips. At the same time, this phylogenetic coherence is strongest for the leading canonical directions and generally decreases for secondary directions, indicating that the dominant composition–environment associations are structured at deeper phylogenetic scales than the weaker, higher-order associations. Consistent with the results from Pagel's $\lambda$ (SI Fig. S16) and the coherence of environmental canonical directions across taxonomic levels (SI Fig. S13), the decrease in coherence for the soil microbiome is more prominent than the decrease in coherence for the ocean microbiome. The observed $\tau_D$ values fall at approximately sub-genus to genus-level phylogenetic depth based on the calibration in Fig. S15.

#### S4.B. Setting a Baseline for Phylogenetic Distance

To interpret the depth values returned by the adjusted consenTRAIT approach, we first established empirical reference distributions for cophenetic phylogenetic distances at each taxonomic level. Cophenetic distance between two OTUs is defined as the sum of branch lengths along the shortest path connecting them through their most recent common ancestor on the GG2 backbone tree. For each level (Genus, Family, Order, Class, Phylum), all OTU pairs were classified as *intra-group* if they shared the same taxon at that level, and *inter-group* otherwise; kernel density estimates of cophenetic distances were computed separately for each combination of level, pair type, and dataset (Fig. S15).

The adjusted consenTRAIT statistic $\tau_D$ is the mean phylogenetic distance from the root of a consensus clade to the tips within it, approximately half the cophenetic distance between two sister tips within the same clade. The reference distributions in Fig. S15 therefore can translate into an expected range for $\tau_D$, for example, a value consistent with Family-level coherence should fall near half the peak of the intra-Family cophenetic distance distribution.

#### S4.C. Pagel's $\lambda$ Approach

Pagel's $\lambda$ [10] quantifies phylogenetic signal by measuring how strongly trait values covary according to shared evolutionary history. Under a Brownian-motion model of trait evolution, closely related taxa are expected to have similar trait values because they share branches of the phylogeny. A value of $\lambda = 0$ indicates no detectable phylogenetic structure, as expected if trait values were randomly assigned to the tips of the tree, whereas $\lambda = 1$ indicates that the observed pattern of trait similarity matches the Brownian-motion expectation on the phylogeny.

Pagel's $\lambda$ was estimated for each canonical direction $j$ and cross-validation fold $f$ using continuous OTU abundance loadings, $\mathbf{b}_{C,\alpha}^{(j,f)}$, as the quantitative trait associated with the tree from the GreenGenes2 backbone. Pagel's $\lambda$ was estimated by maximum likelihood using the `phylolm` package [15] with the model specification `phylolm(y ~ 1, model = "lambda")`. Results are shown in SI Fig. S16.

---


[1] M. Delgado-Baquerizo, A. M. Oliverio, T. E. Brewer, A. Benavent-González, D. J. Eldridge, R. D. Bardgett, F. T. Maestre, B. K. Singh, and N. Fierer, A global atlas of the dominant bacteria found in soil, Science **359**, 320 (2018).

[2] S. Sunagawa, L. P. Coelho, S. Chaffron, J. R. Kultima, K. Labadie, G. Salazar, B. Djahanschiri, G. Zeller, D. R. Mende, A. Alberti, F. M. Cornejo-Castillo, P. I. Costea, C. Cruaud,

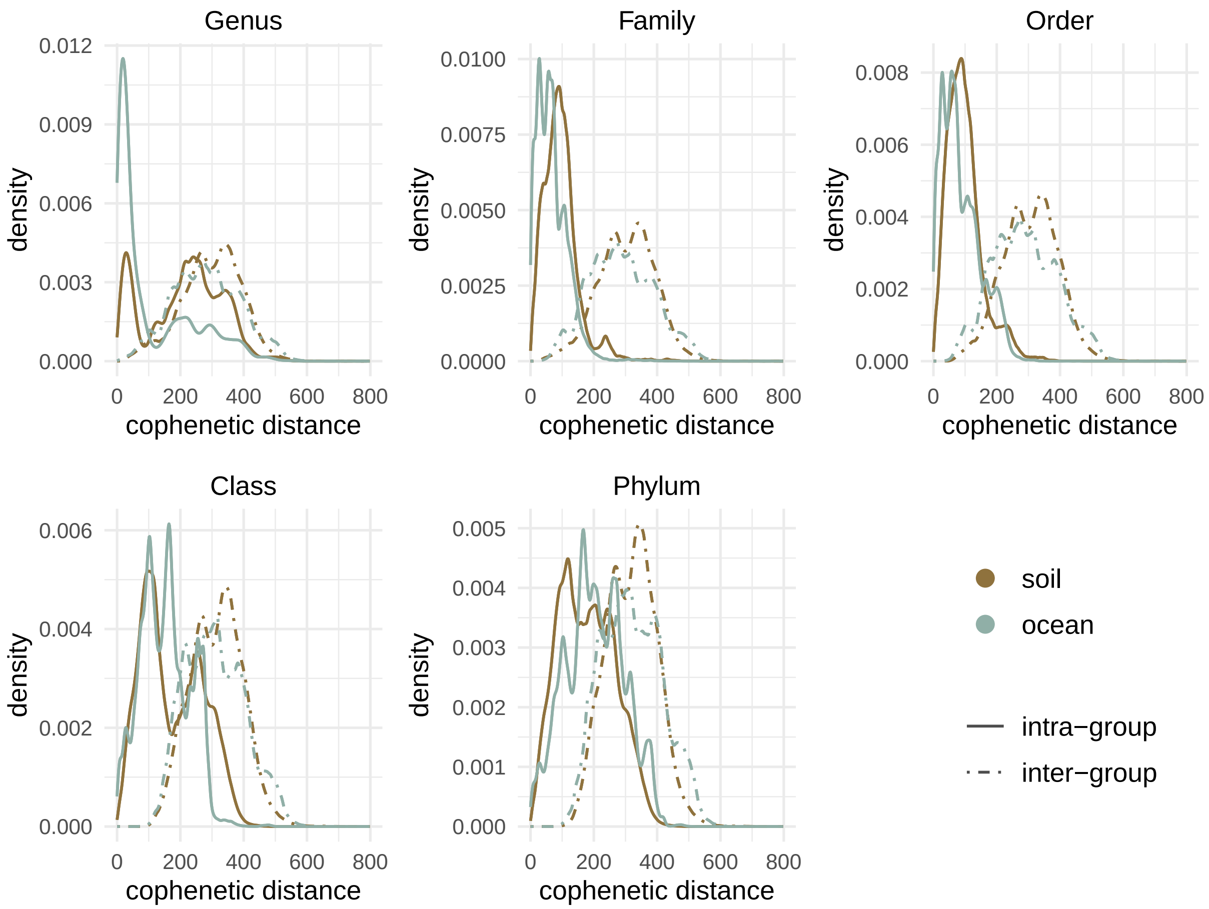


FIG. S15. **Reference distributions of cophenetic distances across taxonomic levels.** Kernel density estimates of pairwise cophenetic distances for OTU pairs that share the same taxon at a given level (*intra-group*, solid lines) and for pairs from different taxa at that level (*inter-group*, dot-dash lines), shown for the soil (brown) and ocean (teal) datasets. Each panel corresponds to one taxonomic level (Genus through Phylum). The $\tau_D$ values reported by the adjusted consenTRAIT approach approximate half the corresponding cophenetic distance; these distributions therefore provide an empirical scale for interpreting $\tau_D$ in terms of taxonomic depth.

F. d'Ovidio, S. Engelen, I. Ferrera, J. M. Gasol, L. Guidi, F. Hildebrand, F. Kokoszka, C. Lepoivre, G. Lima-Mendez, J. Poulain, B. T. Poulos, M. Royo-Llonch, H. Sarmento, S. Vieira-Silva, C. Dimier, M. Picheral, S. Searson, S. Kandels-Lewis, Tara Oceans coordinators, C. Bowler, C. de Vargas, G. Gorsky, N. Grimsley, P. Hingamp, D. Iudicone, O. Jaillon, F. Not, H. Ogata, S. Pesant, S. Speich, L. Stemmann, M. B. Sullivan, J. Weissenbach,

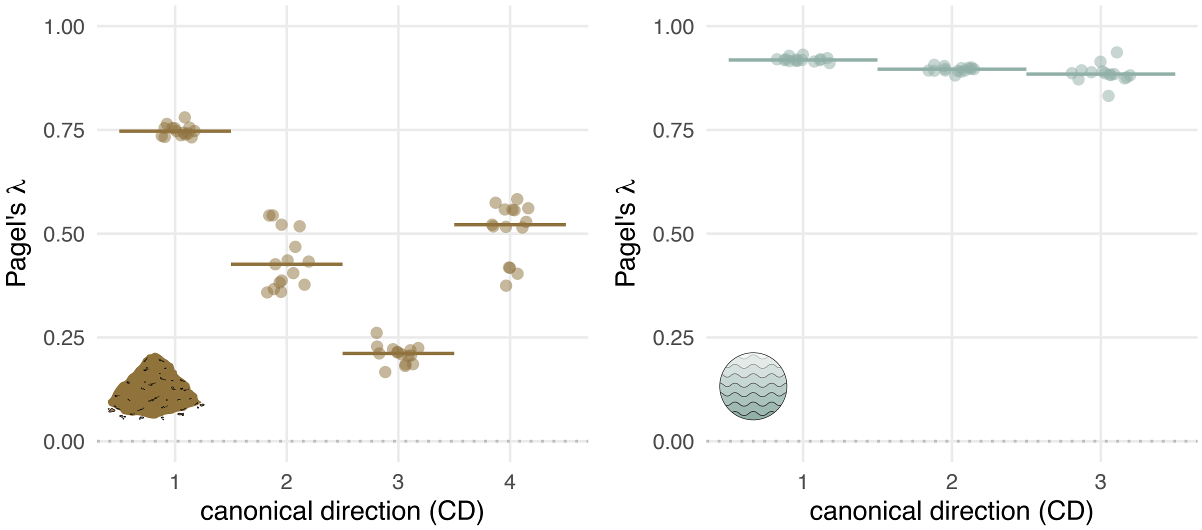


FIG. S16. **Pagel's $\lambda$ decreases from primary to secondary canonical directions.** Observed $\lambda$ values are shown for each significant canonical direction across cross-validation folds; horizontal bars show the median across folds. Colored points show observed values (brown: soil; teal: ocean). Grey points and error bars show the mean ± SD of the null distribution from shuffled loadings, computed separately for each canonical direction and fold index.

P. Wincker, E. Karsenti, J. Raes, S. G. Acinas, and P. Bork, Structure and function of the global ocean microbiome, Science **348**, 1261359 (2015).

[3] R. Logares, S. Sunagawa, G. Salazar, F. M. Cornejo-Castillo, I. Ferrera, H. Sarmento, P. Hingamp, H. Ogata, C. de Vargas, G. Lima-Mendez, J. Raes, J. Poulain, O. Jaillon, P. Wincker, S. Kandels-Lewis, E. Karsenti, P. Bork, and S. G. Acinas, Metagenomic 16S rDNA Illumina tags are a powerful alternative to amplicon sequencing to explore diversity and structure of microbial communities, Environmental Microbiology **16**, 2659 (2014).

[4] L. Guidi, J.-P. Gattuso, S. Pesant, C. Tara Oceans Consortium, and P. Tara Oceans Expedition, Environmental context of all samples from the Tara Oceans Expedition (2009-2013), about carbonate chemistry in the targeted environmental feature (2017).

[5] B. J. Callahan, P. J. McMurdie, M. J. Rosen, A. W. Han, A. J. A. Johnson, and S. P. Holmes, DADA2: High-resolution sample inference from Illumina amplicon data, Nature Methods **13**, 581 (2016).

[6] D. McDonald, Y. Jiang, M. Balaban, K. Cantrell, Q. Zhu, A. Gonzalez, J. T. Morton,

G. Nicolaou, D. H. Parks, S. M. Karst, M. Albertsen, P. Hugenholtz, T. DeSantis, S. J. Song, A. Bartko, A. S. Havulinna, P. Jousilahti, S. Cheng, M. Inouye, T. Niiranen, M. Jain, V. Salomaa, L. Lahti, S. Mirarab, and R. Knight, Greengenes2 unifies microbial data in a single reference tree, Nature Biotechnology **42**, 715 (2024).

[13] M. S. Schmitt, K. K. Lee, F. Bunbury, J. A. Landsittel, V. Vitelli, and S. Kuehn, Learning functional groups in complex microbiomes (2026).

[14] M. Xia, K. Isobe, and J. B. Martiny, Phylogenetic conservation of bacterial environmental responses predicts soil bacterial biogeographic patterns (2026).

[15] L. si Tung Ho and C. Ané, A Linear-Time Algorithm for Gaussian and Non-Gaussian Trait Evolution Models, Systematic Biology **63**, 397 (2014).
